# Rhythmic gene expression and behavioral plasticity in harvester and carpenter ants

**DOI:** 10.64898/2026.04.08.717309

**Authors:** Biplabendu Das, Deborah M. Gordon

**Author notes:** corresponding author to whom proofs will be sent, Email ID of corresponding author.

## Abstract

We examined the overlap in the genes associated with daily rhythms and with behavioral plasticity in ants. We first investigated the daily rhythms of gene expression in the harvester ant, *Pogonomyrmex barbatus*, and how the rhythmic genes overlap with others previously shown to be associated with plasticity of foraging behavior. Then, to consider whether the overlap is conserved across ant species, we compared rhythms of gene expression in the diurnal, desert harvester ants with those previously reported for a distantly related nocturnal, subtropical carpenter ant, *Camponotus floridanus*. First, daily transcriptomes in *P. barbatus* showed that most genes were expressed in light-dark (LD) and constantly dark (DD) conditions at about the same levels; only 11 genes showed at least a two-fold change in expression. Network analysis identified eleven modules of *P. barbatus* genes under LD conditions. Of these 11 clusters, modules C1 and C2 seem to be central nodes of the gene expression network, because they are the most highly connected in LD, and show the strongest preservation in DD vs. LD, and contain core clock gene *Period*. Only one module, C2, showed significant overlap with *P. barbatus* genes that have 24h-rhythmic expression in both LD and DD. There was significant overlap between modules C1, C2, C10, C11, and *P. barbatus* genes found previously to be associated with plasticity in the regulation of foraging activity to manage water loss. A comparison of the daily transcriptome of *P. barbatus* with that of *C. floridanus* showed significant overlap of 24h-rhythmic genes in LD. Modules C1 and C2 of *P. barbatus* also overlap with *C. floridanus* genes previously shown to differ in expression rhythms in nurses and foragers. In combination, these results indicate that genes linking plasticity of the circadian clock and of behavior may be broadly conserved in ants.

## INTRODUCTION

In animals, many behavioral and physiological processes depend on 24-hour cycles of gene expression, regulated by a circadian clock that uses a highly conserved Transcription-Translation-Feedback-Loop (TTFL) (Andreani et al., 2015; Dunlap, 1999; Zhang & Emery, 2012). These cycles of gene expression are flexible in many ways, including in phase, the timing of daily peaks; in amplitude, the range over which daily expression fluctuates; and in periodicity, the interval over which expression repeats. Circadian processes vary with age (Horne & Ostberg, 1976; Roenneberg et al., 2003; Weinert, 2000), social environment (Maury et al., 2020), pathogen infection (Das et al., 2023), seasonal shifts in temperature and light cycles (Gil & Park, 2019; van der Vinne et al., 2014), and with behavioral syndromes in humans (J. Z. Li et al., 2013) (Johansson *et al*., 2016) (Takaesu, 2018).

Rhythmic patterns of behavior, and of gene expression, are entrained by external cues or Zeitgebers (Saunders, 2002). In social insects, entrainment depends on external cues such as light and temperature (Giannoni-Guzmán et al., 2021; Ingram et al., 2012; Moore, 2001; Roces & Núñez, 1996), but also on social cues, such as volatile pheromones and substrate vibration (Siehler & Bloch, 2020). Many cues can operate together to entrain rhythmic cycles. For example, ants kept in darkness can synchronize to light-dark (LD) cycles outside the nest using social interactions with nestmates that have access to LD cycles (Lone & Sharma, 2011).

In constant darkness (DD) and in the absence of external rhythmic cues, the expression of some genes, and some behavioral processes, show near 24h-rhythmic patterns of expression (Saunders, 2002). The period of the free-running clock is usually slightly lower or higher than 24h, which causes the entire rhythm to shift over time (Aschoff, 1981). Here we use the term “core clock genes” to refer to the components of the highly conserved transcription-translation feedback loop that drives near-24h rhythm in animals, including *Period* (*Per*) and *Clock*, while we use “clock-controlled genes” to refer to those whose expression shows 24h rhythm in DD. Similar definitions have been used in other studies (Lech et al., 2016).

Clock-controlled genes that show 24h-rhythmic expression in both LD and DD can show a phase shift in DD, free-running conditions, after a few days in the absence of external 24h rhythms. For example, the phase of the 24h-rhythmic *foraging* (*For*) gene expression shows a shift in constant dark vs. LD conditions in ants, including *Solenopsis invicta* (Lei *et al*., 2019) and *Pogonomyrmex barbatus* (Ingram *et al*., 2016); so does the clock gene *Per* in *P. occidentalis* (Ingram *et al*., 2009). Free-running conditions also influence ant foraging rhythms (Mildner & Roces, 2017; Sharma *et al*., 2004).

Behavioral plasticity is associated with plasticity of rhythmic gene expression. In social insects, task groups differ in their gene expression profile: in ants (Ingram et al., 2009, 2016; Manfredini et al., 2014; Mikheyev & Linksvayer, 2015; Qiu et al., 2017), honeybees (Bloch et al., 2004) and other social insects (Araujo & Arias, 2021; Huisken & Rehan, 2023; Saiki et al., 2022; Vleurinck et al., 2016). These task-specific patterns of gene expression are correlated with task-specific activity rhythms. In honeybees, brood care workers and foragers differ in daily expression of several key clock genes (Rodriguez-Zas et al., 2012). In the carpenter ant *Camponotus floridanus,* brood care workers and foragers differ in the periodicity of the core clock gene *Per* expression (Das & de Bekker, 2022), and in many other genes probably associated with *Per*. In *P. occidentalis* (Ingram et al., 2011), task groups differ in the daily periodicity of *Per*, and its phase shifts seasonally (Ingram et al., 2009).

In this study, we examine how clock genes are linked to behavioral plasticity in two distantly-related and ecologically distinct ant species, *P. barbatus* and *C. floridanus*, to consider whether these links are broadly conserved across ants. We first characterized the daily expression profiles of all genes, including the core clock gene *Per*, of the harvester ant *P. barbatus* in LD and DD conditions. We examined gene expression in the brains of *P. barbatus,* using newly sequenced time-course RNASeq data along with previous RNASeq data linked to behavioral plasticity (Friedman et al., 2020). Then, to consider whether these links are broadly conserved across ants, we compared the results from the diurnal, desert harvester ant *P. barbatus*, with patterns previously found in the distantly related, nocturnal, subtropical carpenter ant, *C. floridanus* (Das & de Bekker, 2022).

Our study has 4 sections: 1) Comparison of gene expression in DD and LD in harvester ants, 2) Comparison of gene co-expression networks in DD vs. LD in harvester ants, 3) Overlap between genes related to clock-control and those regulating foraging behavior in harvester ants, and 4) Comparison of rhythmically expressed genes that regulate task behavior in harvester ants and carpenter ants.

In sections 1 and 2, we compare the entire daily transcriptome of *P. barbatus* ant brains in LD and DD to characterize the genes that show 24h-rhythmic expression and to identify clock-controlled genes. This extends previous work linking plasticity of the circadian clock and behavioral plasticity in ants, which have focused on a few genes instead of the entire transcriptome. We draw on methods that model daily transcriptomic data into a co-expression network, using Weighted Gene Co-expression Network Analysis (WGCNA) (Langfelder & Horvath, 2008), This method uses clustering techniques and network statistics to identify gene clusters or modules that show similar temporal patterns of expression; genes in a module are most likely co-expressed in time.

WGCNA has been used in comparative transcriptomics studies of time-series RNASeq data (Pal et al., 2025; Wang et al., 2022; Zacharias et al., 2025), because it circumvents the limitations of a direct comparison. Which genes are classified as significantly 24h-rhythmic depends on the choice of a rhythmicity-detecting algorithm and the use of an arbitrary p-value cutoff (Laloum & Robinson-Rechavi, 2020), creating disparities in the gene sets identified across different studies and species (Morandin et al., 2017). Here we avoid this problem by using unsupervised clustering methods to identify gene clusters that are highly co-expressed in time as a reference network. If two gene sets of interest overlap with the same gene module, we infer a link between the two gene sets and the biological process it regulates.

In section 3, we examine the overlap between clock-controlled genes of *P. barbatus* that show 24h-rhythmic in DD, and the genes associated with behavioral plasticity, in particular the regulation of foraging activity to manage water loss (Friedman *et al*., 2020). Foragers lose water to evaporation while out searching for the seeds they eat. Some colonies reduce foraging activity on dry days, sacrificing food intake to conserve water, while other colonies do not (Gordon, 2013; Gordon et al., 2023). Colonies that differ in the regulation of foraging show differences in gene expression, including in genes associated with dopamine neurophysiology (Friedman *et al*., 2018, 2020). We examined the overlap between these genes identified in previous work, and the clock-controlled genes we identified here.

In section 4, we compare the overlap between the clock-controlled *P. barbatus* genes involved in the regulation of foraging behavior and those in *C. floridanus* involved in task plasticity from brood care workers to foragers (Das & de Bekker, 2022). These two species are phylogenetically distantly related, from different subfamilies (Myrmicinae and Formicinae) that diverged around 100 million years ago (Nelsen et al., 2018). The two species differ in behavioral rhythms and foraging behavior. The desert harvester ant is a diurnal species (Gordon, 1984), with large colonies (Gordon, 1992) of monomorphic workers.

Foragers search for scattered seeds in a daily temporal pattern that avoids the hot dry conditions at midday (Pagliara et al., 2018). In contrast, the omnivorous species *C. floridanus* is native to temperate regions, with large colonies of polymorphic workers that are mostly nocturnal, with daily peaks of foraging around dusk and dawn (Das & de Bekker, 2022; Feldhaar et al., 2007; Kay et al., 2018). Shared patterns of gene expression in these two distantly related species would suggest that the patterns are broadly conserved across ants.

## MATERIALS AND METHODS

Gene expression was measured in ants from a queenright laboratory colony of the red harvester ant *Pogonomyrmex barbatus*, with brood and about 300 workers, collected in August 2022 near Rodeo, New Mexico.

### Experimental setup and sampling methods for sections 1-2

#### Colony setup in the lab

The colony was maintained at constant temperature in two dark, enclosed nest boxes connected in series to an open foraging arena by clear, vinyl tubes of 5/16“ inner diameter. The dimensions of the foraging arena were 4 x 3 feet, while the nest boxes were 7 x 3 inch. The floors of the nest boxes were made of plaster and watered at least two times a week to maintain humidity. Food (crickets and sucrose solution) and water were provided once a week in the foraging arena. The queen and the brood stayed in the nest box farthest from the foraging arena.

The foraging arena was first exposed to oscillating cycles of 12h of light and 12h of dark (12:12 LD cycle) for 7 days to allow the ants to entrain or synchronize their internal rhythms to the LD cycle. The temperature, humidity, and light levels experienced by the colony during the 7 days of the entrainment period are shown in Supplementary Fig. 1. After 7 days of entrainment to 12:12 LD cycle, the foraging arena was kept in DD for 3 days to allow the internal clock of the ants to free run.

#### Monitoring colony activity rhythm

We monitored the activity of the ants in the transparent tube, connecting the outer dark nest box to the foraging arena, which was exposed to the LD cycles using timelapse videos that captured a photo every 30 seconds continuously. Using the videos, we counted the number of ants observed in the tube in 6 consecutive frames, each 30 seconds apart, over a 3-minute window, in each hour during the daytime (Supplementary Fig. S2). Total activity for each hour was the sum of the 6 counts.

We could not record video in the dark during LD, so our video-recordings tracked activity patterns only during the 12h of daytime, with lights on. Instead, we used an indirect method. We determined whether there were repeated patterns with a periodicity of 12 hours or less, by stitching together the 12h day-time activity profiles over 7 days to create a continuous timeseries. We used rhythmicity in day-time activity as a proxy to confirm entrainment to LD.

To confirm rhythmicity of the daytime activity, we performed wavelet decomposition and periodicity detection using the WaveletComp package (Roesch et al., 2014; Rösch & Schmidbauer, 2018) (Supplementary Fig. S3). We looked for waveforms (Fourier periods) in the concatenated timeseries with periodicity between 2h and 12h, using the “analyze.wavelet” function that produces a wavelet power spectrum (Supplementary Fig. S3A). The power of each periodicity between 2h and 12h is a measure of how much it contributes to the observed fluctuations.

Significant periodicities were detected using the “wt.avg” function on the wavelet power spectrum; significance was inferred when p < 0.01 (red dots in Supplementary Fig. S3B).

#### Sampling design and collection method

We obtained daily transcriptomes of ant brain tissues under two conditions: (1) exposed to 12:12 LD cycle for 7 days, after entrainment was confirmed, and (2) exposed to 24h DD for three consecutive days following the LD sampling (Supplementary Fig. 1A).

For both LD and DD conditions, we collected three ants every 2h, over a single 24h period. We refer to the timepoints for the LD experiment as Zeitgeber Time or ZT, and for the DD experiment as Circadian Time or CT. At each timepoint, we pooled three ant brains to perform a single sequencing run. We then used mean expression across the three ant brains and did not measure the variance.

Ants were collected from the foraging arena during the actual or subjective daytime (1h-11h ZT or CT, respectively). Ants that go outside the nest usually stay close to the entrance when not active (Gordon et al., 2005). During night time (13h-23h ZT or CT, respectively), we collected ants in the tube linking the foraging arena and nest box, close to the entrance to the arena, as in (Das & de Bekker, 2022).

For the LD experiment, we sampled foragers every 2h starting at ZT7 on Experimental Day 8 and ending at ZT5 on Day 9 (Supplementary Fig. 1A). Next, the colony was released to DD for 3 consecutive days, and then foragers were sampled in DD every 2h starting at CT7 on Day 13 and ending at CT5 on Day 14 (Supplementary Fig. 1A). During LD cycles, lights turned on at ZT0 (same as ZT24) and turned off at ZT12.

### RNA extraction, library preparation and RNASeq

After collection, each ant was individually placed in a cryotube, immediately flash frozen in liquid nitrogen, and stored at-80°C. Prior to RNA extractions, we dissected and pooled the brains from the three ants collected at each time point, for a given light-dark cycle, in a frozen cryotube containing three ball bearings and 100 µL of TRIzol (Invitrogen). Next, we used a Bead Mill 24 (Fisherbrand) to homogenize the pooled brain tissues at 4.5 m/s for 60 secs. We extracted total RNA from the homogenized brain tissues of ants and prepared cDNA libraries from extracted mRNA. For each cDNA library, we extracted mRNA with poly-A magnetic beads (NEB) from 500 ng of total RNA, which was converted to 270-330 bp cDNA fragments using the Ultra II Directional Kit (NEB). To enable sequencing different samples on the same lane, we added unique adapters (NEB) to each cDNA library. Supplementary Figure 5 shows the concentrations of extracted mRNA and the resulting cDNA libraries sequenced for each timepoint.

All twenty-four cDNA libraries were sequenced as 150 bp paired-end reads on a NovaSeq 6000. In total, the sequencing resulted in a mean of 23 million reads (± 2 million, SD) per sample, with a minimum of 19 and a maximum of 28 million reads. To process the sequenced transcriptomes, we first removed sequencing adapters and low-quality reads with BBDuk (Bushnell, 2018), and then used HISAT2 (Kim et al., 2017) to map transcripts to the latest *P. barbatus* genome, Pbar_UMD_V03 (Smith *et al*., 2011). We then normalized the expression of each gene, for a given sample, to Fragments Per Kilobase of transcript per Million (FPKM) using Cuffdiff (Trapnell *et al*., 2012).

### Data analyses

Data analysis methods are summarized in Table 1. Data analyses were performed in RStudio (Team, RStudio, 2016) using the R programming language v3.5.1 (Team, R Core, 2013). Heatmaps were generated using the pheatmap (Kolde, 2019) and viridis (Garnier *et al*., 2018) packages. Upset diagrams used to visualize intersecting gene sets were generated using the UpsetR package (Conway et al., 2017), and all other figures were generated using the ggplot2 package (Wickham, 2011).

**Table 1.**
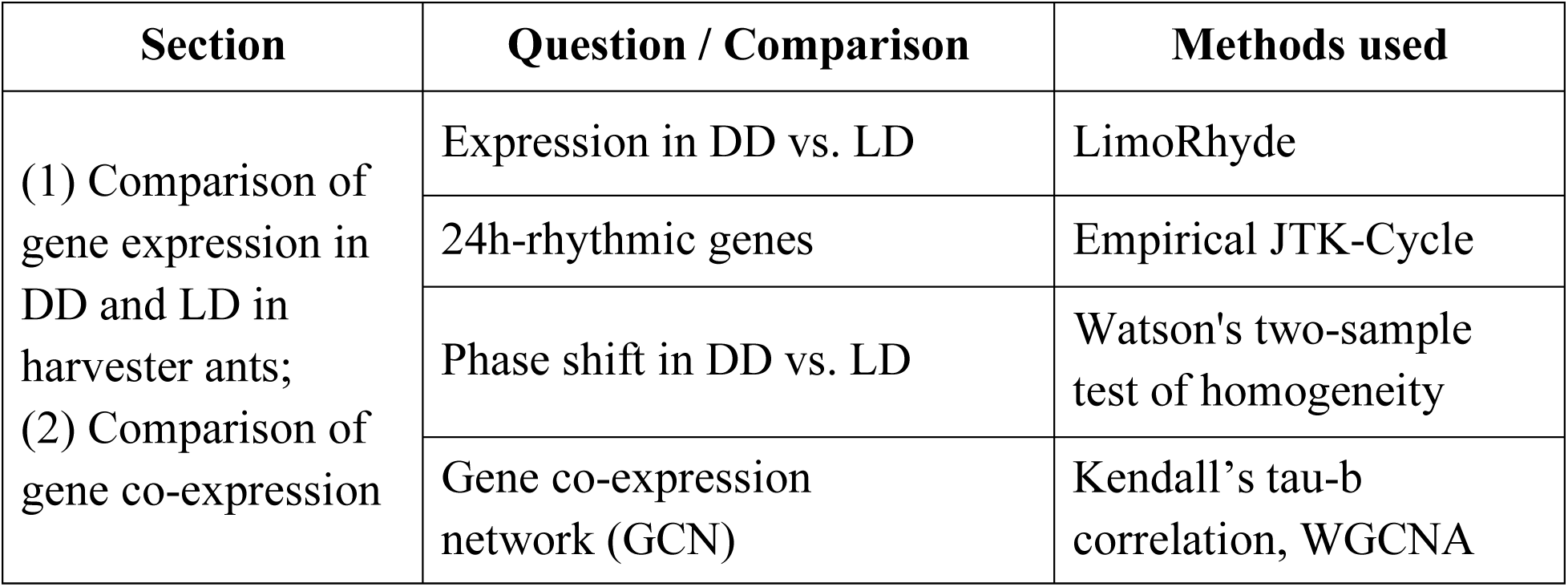
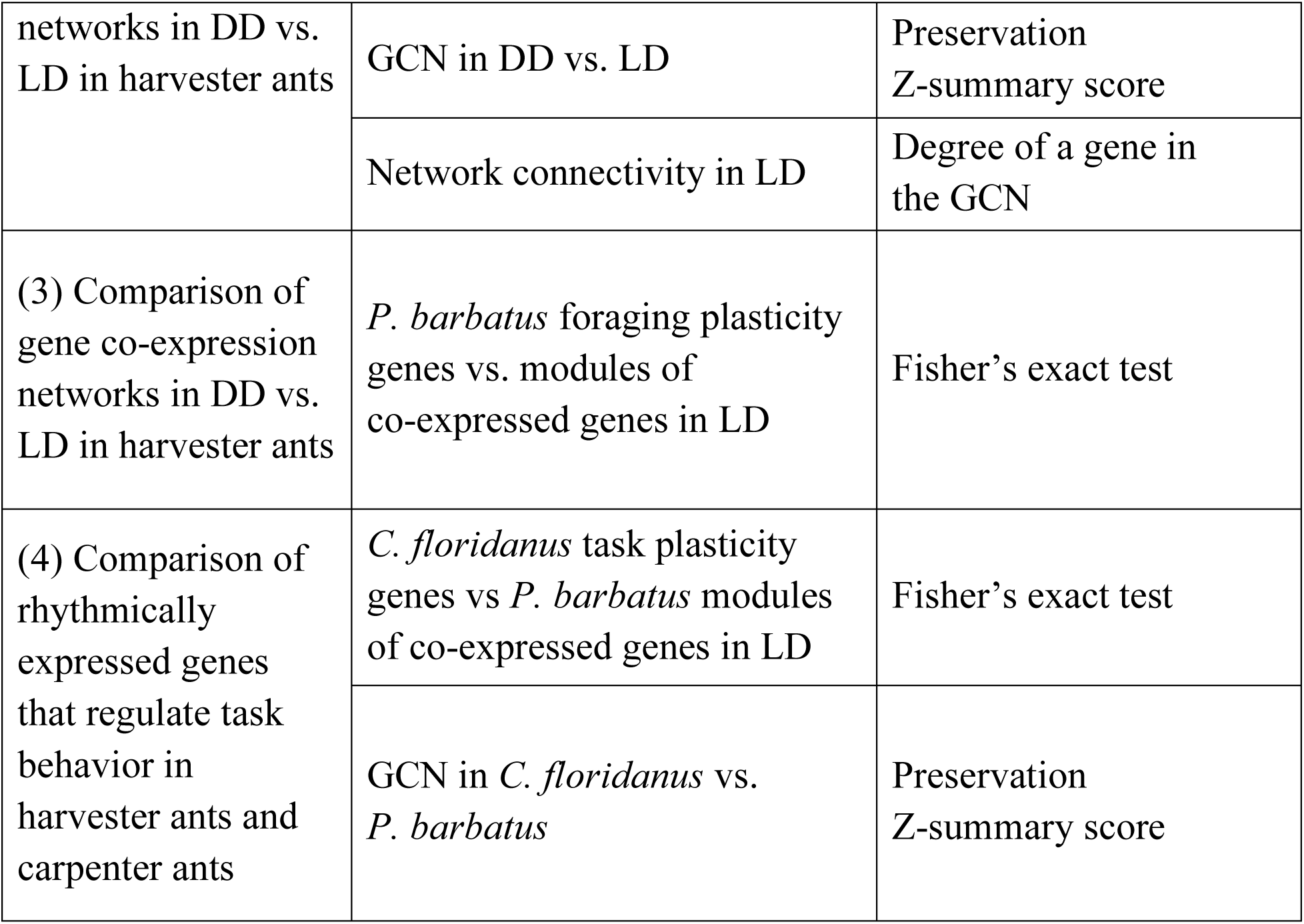
Summary of data analysis methods.

### A. Identifying protein domains overrepresented in a set of genes

For a given set of genes, we quantified the overrepresented Pfam terms (protein domains) in the gene set using a hypergeometric test via the “check_enrichment” function in the timecourseRnaseq package on GitHub (https://github.com/biplabendu/timecourseRnaseq) (Das, 2022). Significance was assessed at 5% FDR. We used the entire genome (all genes in *P. barbatus*) as the background geneset for identifying overrepresented Pfam terms in the set of genes that are not expressed in the brains. For all other sets of *P. barbatus* genes, unless specified otherwise, we used all expressed genes (≥ 1 FPKM expression for at least one sample) as the background geneset for identifying overrepresented Pfam terms.

We tested for overrepresentation only for Pfam terms annotated for at least 5 protein coding genes of *P. barbatus*. We obtained the Pfam annotations for the *P. barbatus* genome, Pbar_UMD_V03 (Smith *et al*., 2011) using Interproscan (Jones *et al*., 2014).

### B. Modeling a gene co-expression network

We modelled the transcriptomic dataset as a GCN to compare changes in the network properties of the transcriptome in free-running DD conditions relative to LD. Modelling was performed using clustering functions from the WGCNA package (Langfelder et al., 2011; Langfelder & Horvath, 2008, 2014). Gene-gene and module-module correlations for similar expression over the 24h day were calculated using Kendall’s tau-b correlation (Samara & Randles, 1988), and the global connectivity patterns of the gene co-expression network (GCN) were visualized using the igraph package (Csardi, 2013).

### C. Identifying overlaps between different gene sets of interest

To identify links between rhythmically expressed genes and those previously associated with behavioral plasticity, we used a network-based approach with the following steps:

(a) construct a model of gene co-expression network (GCN) using the time-course RNASeq data for all expressed genes in LD, which identifies clusters or modules of genes that show strong co-expression in time; (b) annotate the GCN to identify the modules in the network that show an overlap with the gene sets of interest, and

(b) determine whether multiple gene sets of interest show any overlap with the modules that contain most of the rhythmically expressed genes.

Significant overlap between a given module and a gene set of interest was assessed by Fisher’s exact tests using the GeneOverlap package (Shen, 2016). Benjamini-Hochberg corrections were applied to correct p-values for multiple comparisons.

### 1. Comparison of gene expression in DD and LD in harvester ants

#### Differentially expressed genes in DD vs. LD

To determine whether genes were differentially expressed in free-running DD conditions, compared to LD, we used the linear modelling framework of LimoRhyde (Singer & Hughey, 2019). A gene was considered to be differentially expressed if treatment was a significant predictor, at a 5% false discovery rate (FDR), and the gene showed at least a two-fold change in mean diel expression between LD and DD conditions (abs(log_2_-fold-change) ≥ 1).

#### 24h-rhythmic genes

To test for significant 24h rhythms in gene expression, we used the rhythmicity detection algorithm, empirical JTK-Cycle (eJTK) (Hutchison et al., 2015, 2018). Only genes that had diel expression values of FPKM ≥ 1 for at least half of all sampled timepoints were tested for rhythmicity. A gene was considered to be significantly 24h-rhythmic if it had a Gamma p-value < 0.05. Once we identified the *P. barbatus* genes that oscillate in LD and DD conditions, we identified the location of these genes in the *P. barbatus* GCN.

#### Phase shift of 24h-rhythmic genes

To test whether the *P. barbatus* genes expressed with a 24h rhythm show a phase shift in DD vs. LD, we first obtained the estimated phase of all genes that show 24h-rhythmicity in LD and DD from the output of eJTK. We then converted the phases to polar coordinates using the circular package for R and ran the Watson test for the two circular distributions using the ‘watson.two.test’ function in the stats package for R (Team, R Core, 2013).

We also visually assessed whether a phase shift occurred in the set of genes expressed with a 24h-rhythm in both LD and DD. We clustered the 345 genes showing 24h-rhythmic gene expression in both LD and DD according to their daily expression using an agglomerative hierarchical clustering framework (method = complete linkage) via the ‘hclust’ function in the stats package for R (Team, R Core, 2013), and visualized it as a heatmap using the pheatmap package (Kolde, 2019).

### 2. Comparison of gene co-expression network in DD vs. LD in harvester ants

To characterize how the network of gene expression changes in constant dark conditions, relative to LD, we first modelled the daily transcriptome of forager brains in LD as a co-expression network. Then, using the gene co-expression network in LD as the reference, we identified the changes in the daily transcriptome of forager brains in DD conditions, using the protocol described in (Langfelder *et al*., 2011).

#### Network connectivity

We compared the average total connectivity of the modules of *P. barbatus* genes in DD, compared to LD. For each gene, the total connectivity (or degree) of a gene is defined as the sum of connection strengths with the other genes in the network (Langfelder & Horvath, 2008). A module containing genes with high average connectivity indicates that those genes are tightly co-expressed in time.

#### Network Preservation

We asked whether the co-expressed gene modules identified in LD showed the same connectivity patterns in DD by comparing the Preservation Zsummary score described in (Langfelder *et al*., 2011). A Zsummary > 10 indicates strong evidence that the structure of the module is preserved, 2 < Zsummary < 10 indicates moderate evidence, and Zsummary < 2 provides no evidence for module preservation (Langfelder *et al*., 2011).

### 3. Overlap between gene co-expression networks in DD vs. LD in harvester ants

We examined whether the *P. barbatus* clock-controlled genes, which show 24h-rhythmic expression in DD (*rhy24-Pbar-DD*), were the same as those associated in previous work with reducing foraging activity in dry conditions (Friedman *et al*., 2018, 2020; Gordon *et al*., 2023). We obtained the top 5% (516 genes) of all genes that in previous work were associated with reducing foraging in dry conditions (*Pbar-rH-pos*) or with not reducing foraging in dry conditions (*Pbar-rH-neg*) (Friedman *et al*., 2020).

We examined the overlap of the two sets of *P. barbatus* genes, *Pbar-rH-pos* and *Pbar-rH-neg,* whose expression is associated with the regulation of foraging behavior, with the 11 modules of co-expressed genes identified in the *P. barbatus* brain GCN under LD conditions. Significant overlap between sets of genes was measured using Fisher’s exact test; significance was inferred when p < 0.05.

### 4. Comparison of rhythmically expressed genes that regulate task behavior in harvester ants and carpenter ants

To compare daily patterns of gene expression in *Camponotus floridanus* and *P. barbatus*, we identified the one-to-one orthologous genes between the two ant species using proteinortho5 (Lechner *et al*., 2011) (Data S1). For time-course transcriptomes of *C. floridanus*, we used data from (Das & de Bekker, 2022) which employed the same protocols, including the LD entrainment conditions and sampling frequency, as the present study. All cross-species analyses were performed by converting *C. floridanus* gene names to their *P. barbatus* orthologs.

First we tested whether the set of genes with 24h-rhythmic expression in ant brains, under LD conditions, overlapped between *C. floridanus* (*for24-Cflo-LD*) (Das & de Bekker, 2022) and *P. barbatus* (*for24-Pbar-LD*).

Next, we asked whether some of the modules of co-expressed genes identified in the diurnal *P. barbatus* brains, under LD, show preservation of their connectivity patterns in the distantly related and nocturnal *C. floridanus* brains under the same light-dark conditions. We compared the Preservation Zsummary score as described above, with the model of *P. barbatus* GCN in LD conditions as the reference network, and the normalized gene expression data for *C. floridanus* forager brains in LD condition (Das & de Bekker, 2022) as the comparison group.

Finally, to evaluate the similarity of the clock genes associated with the regulation of foraging in *P. barbatus* and with task in *C. floridanus*, we compared the overlap between previously reported *C. floridanus* genes that show task-associated differentially rhythmicity (*for24nur8-Cflo*) with the 11 modules of co-expressed genes identified here in the *P. barbatu*s brain GCN. The set, *for24nur8-Cflo*, represent genes whose expression oscillates with a 24h period in *C. floridanus* forager brains but with a 8h period in nurse brains (Das & de Bekker, 2022).

## RESULTS

### 1. Comparison of gene expression in DD and LD in harvester ants

Most genes were expressed, FPKM ≥ 1 for at least one of the 12 sampled time points, in both LD and DD. Of the 12557 genes currently annotated in the *P. barbatus* genome (Smith *et al*., 2011), 87% (10906 genes) were expressed during the 12:12 LD condition, 84% (10537 genes) were expressed during the DD condition, and 83% (10446 genes) were expressed in both LD and DD conditions (Data S2). There was significant overlap between the two gene sets (Fisher’s exact test; odds-ratio = 389, p < 2.2e-16).

#### Differentially expressed genes in DD vs. LD

The genes expressed in both LD and DD were expressed at similar levels in both conditions (fold-change < 2 at 5% FDR; Fig. 1A). Only ten of the 10446 *P. barbatus* genes expressed in both conditions showed at least a two-fold increase, and a single gene, *trehalose transporter*, showed at least a two-fold decrease, in average daily expression in DD relative to LD (Fig. 1, Data S2).

**Figure 1.**
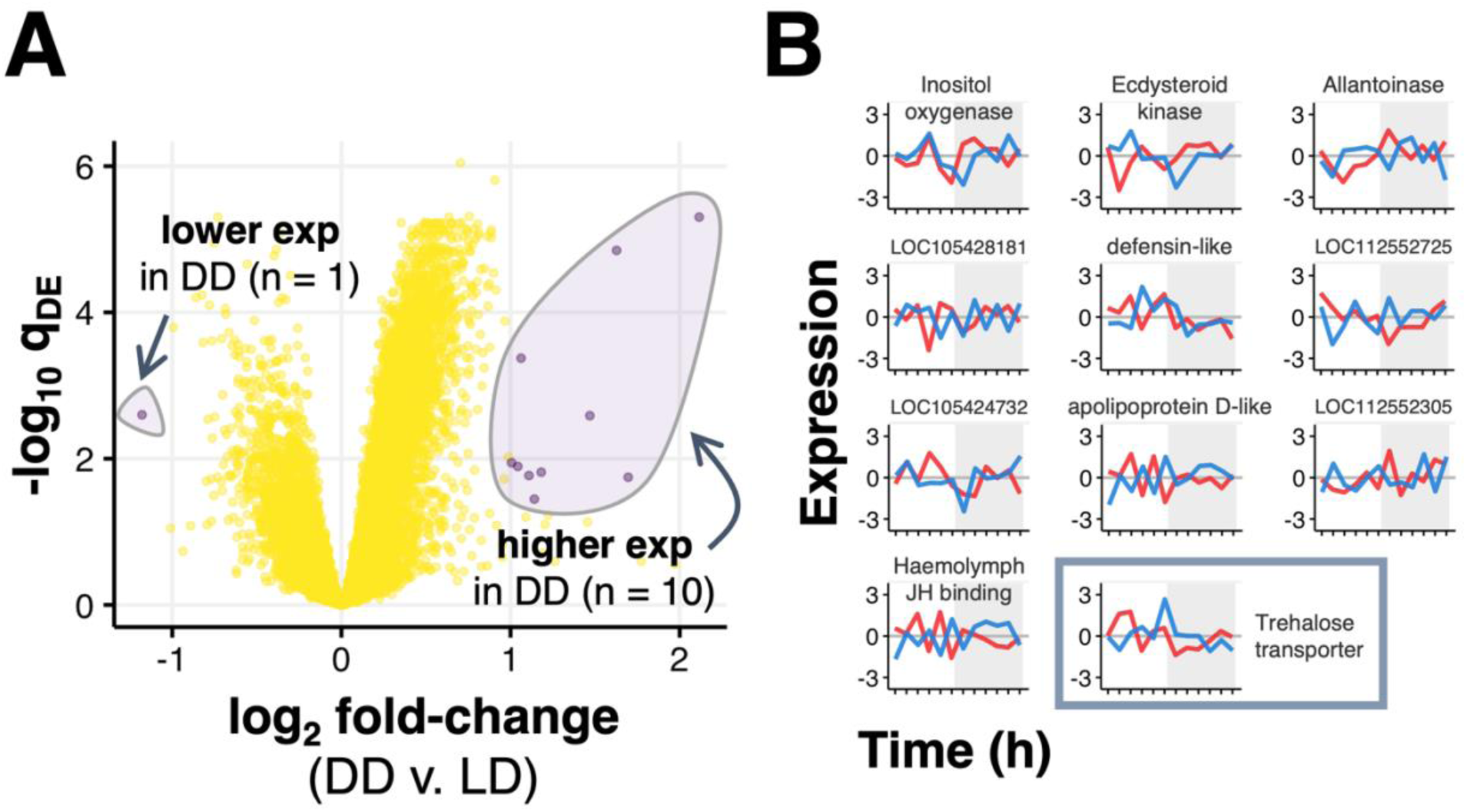
Differences in gene expression in DD and LD conditions. (**A**) The x-axis shows the change (log2-scale) in the expression of a gene in DD vs LD. The y-axis shows the Benjamini-Hochberg corrected p-value (q_DE_); higher y-values represent smaller p-values. Each dot represents one gene. Purple: fold-change ≥ 2 and q_DE_ < 0.05; yellow: not differentially expressed. (**B**) Daily temporal patterns of expression for the 11 genes differentially expressed in DD and LD. Red lines show the daily pattern of gene expression in LD and blue lines in DD. The y-axis shows amount of gene expression (z-scored, log2-transformed FPKM values), and the x-axis shows time in hours (ZT in LD and CT in DD). Trehalose transporter (bottom right) is the only gene downregulated in DD relative to LD. For genes that are not characterized, the gene ID is shown.

Next, we examined the daily expression patterns of the eleven genes differentially expressed in DD compared to LD (Fig. 1B). The gene *Inositol oxygenase*, which undergoes a two-fold increase in average expression levels in DD vs. LD, shows a rhythmic pattern in both LD (red line) and DD (blue line) conditions: a peak around the middle of the daytime and another peak in the middle to late nighttime. The gene *Ecdysteroid kinase*, also expressed more in DD than LD, shows rhythmic patterns of expression with a phase shift in DD vs. LD.

The other 9 genes differentially expressed in DD relative to LD showed no apparent daily rhythms.

#### 24h-rhythmic genes

When we tested for 24h rhythms of daily expression using eJTK cycle (Hutchison *et al*., 2015), we found similar numbers of genes expressed with 24h rhythms in LD and DD conditions, and a smaller set that are 24-h rhythmic in both conditions. A set of 1536 *P. barbatus* genes were classified as significantly 24h-rhythmic in forager brains in LD (*for24-Pbar-LD*), a set of 1547 genes were classified as 24h-rhythmic in DD conditions (*for24-Pbar-DD*), and 345 genes were classified as 24h-rhythmic in both conditions (Data S4). There was significant overlap between the two sets of genes, between *for24-Pbar-LD* and *for24-Pbar-DD* (Fisher’s exact test: overlap: 345, odds-ratio: 2; p < 0.001).

#### Phase shift of 24h-rhythmic genes

Daily gene expression showed changes in phase of daily expression in DD vs. LD conditions. The majority of genes classified as significantly 24h-rhythmic in LD (red) show a daily peak of expression around the middle of the 24h day with peaks mostly at night-time (Supplementary Fig. S4A). In comparison, the majority of genes classified as significantly 24h-rhythmic in DD (blue) show a daily peak of expression at early daytime (Supplementary Fig. S4B).

A phase shift in the daily gene expression was also confirmed for the 345 genes that were classified as 24h-rhythmic in both DD and LD (Watson’s two-sample test of homogeneity: U^2^ = 4.91, p < 0.001) (Fig. 2). The gene expression of these 345 genes in LD and DD conditions is shown in the heatmap in Figure 2, where brighter colors indicate higher expression and darker colors indicate lower expression. In LD conditions, left panel in Figure 2, the majority of genes show peak daily expression between ZT7 and ZT17. The same *P. barbatus* genes in DD, right panel in Figure 2, show 24h-rhythmic, phase-shifted daily expression relative to LD. The peak expression of these genes, which was between ZT7 and ZT17 in LD, now occurs between CT21 and CT5.

**Figure 2.**
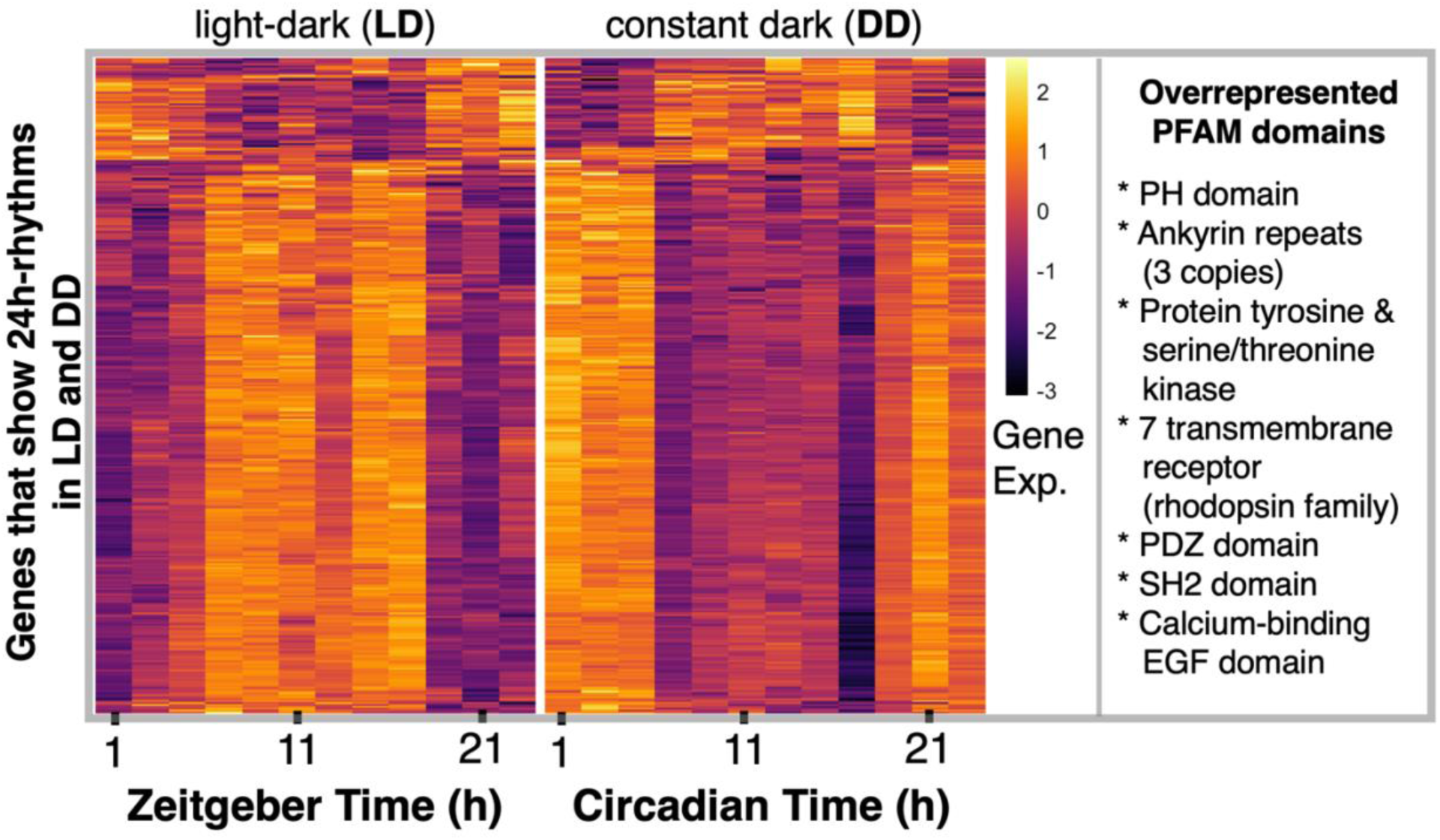
Shifts in 24h-rhythmic gene expression under DD vs LD. Heatmap of daily expression (z-score) patterns of the 345 genes expressed with a 24h-rhythm in *P. barbatus* brains in both LD and DD conditions. Each row represents a single gene, and each column represents the time of day at each 2 hr interval at which a sample was collected, shown in chronological order from left to right. Colors indicate normalized gene expression values (z-scores): negative numbers (dark colors) indicate lower gene expression relative to their daily average, and positive numbers (lighter colors) indicate higher gene expression. The box on the right lists the Pfam domains overrepresented in this set of 345 genes.

Genes that regulate the insect circadian clock, including *Per*, and those associated with behavioral plasticity both showed phase shifts in DD conditions, relative to LD. Figure 3 shows changes in the daily expression patterns of selected genes shown in previous studies to regulate the insect clock, to be associated with behavioral plasticity, or to have the same tempo as *Per*.

**Figure 3.**
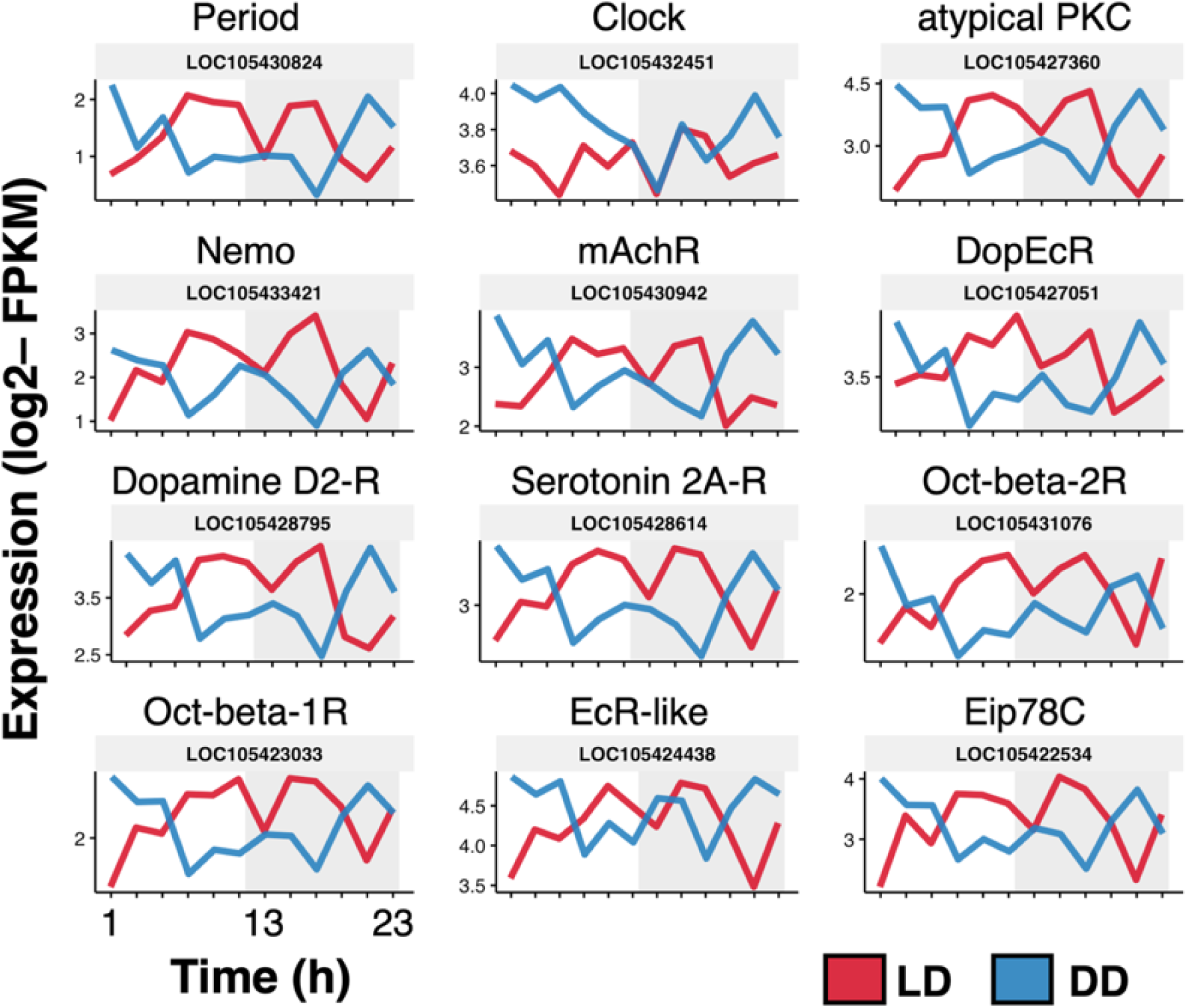
P*eriod*-like phase shifts were observed for several genes linked to the insect circadian clock and behavioral plasticity. The y-axis shows amount of gene expression (log2-transformed FPKM values), and the x-axis shows time in hours (ZT in LD and CT in DD). The shaded portion indicates nighttime in LD (ZT12 to ZT24) and subjective nighttime in DD (CT12 to CT24).

The core clock gene *Per,* one of the 345 *P. barbatus* genes that is significantly 24h-rhythmic in LD and DD, shows a phase shift in DD relative to LD conditions. In LD (red lines in Fig. 3), *Per* shows a daily peak of expression around the middle of the 24h day between ZT7 and ZT17, but in DD (blue lines), its daily peak shifts to the end/beginning of the 24h day between CT21 and CT1 (Fig. 3). The genes with a daily expression pattern similar to *Per* in LD include genes either related to the clock (*Clock, atypical PKC, and Nemo*) or to behavioral plasticity in ants (*mAchR, DopEcR, Dopamine D2-R, Serotonin 2A-R, Oct-beta-1R, Oct-beta-2R*). All show a gradual increase in their expression levels in the early morning, with a first peak around the middle of the daytime (ZT7), a small dip right after lights are turned off (ZT13), and a second peak around the middle of the nighttime (ZT17), followed by a slow decline to a minimum level at night before the lights turn on (ZT21) (Fig 3, red lines).

In DD (Fig 3, blue lines), most of these genes show a phase shift in DD similar to that of *Per* (*atypical PKC, Nemo, mAchR, DopEcR, Dopamine D2-R, Serotonin 2A-R, Oct-beta-1R, Oct-beta-2R*). The pattern is almost opposite to that of LD, with a daily peak of expression during late night or early morning, followed by a slow decline during the daytime, with the first dip around the middle of the subjective daytime (CT7) the second dip around the middle of the subjective nighttime (CT17), and finally a gradual increase during the rest of the subjective nighttime. Finally, *Per-*like expression in LD and DD is not visually evident in other genes that have zig-zag patterns (*EcR-like, Eip78C*) (Fig. 3).

### 2. Comparison of gene co-expression network in DD vs. LD in harvester ants

We modeled the daily transcriptome of *P. barbatus* as a gene co-expression network (GCN) using 9925 genes that were expressed in LD for at least half of all sampled time points, which identified eleven modules or clusters of co-expressed genes (Fig. 4A; Data S3).

**Figure 4.**
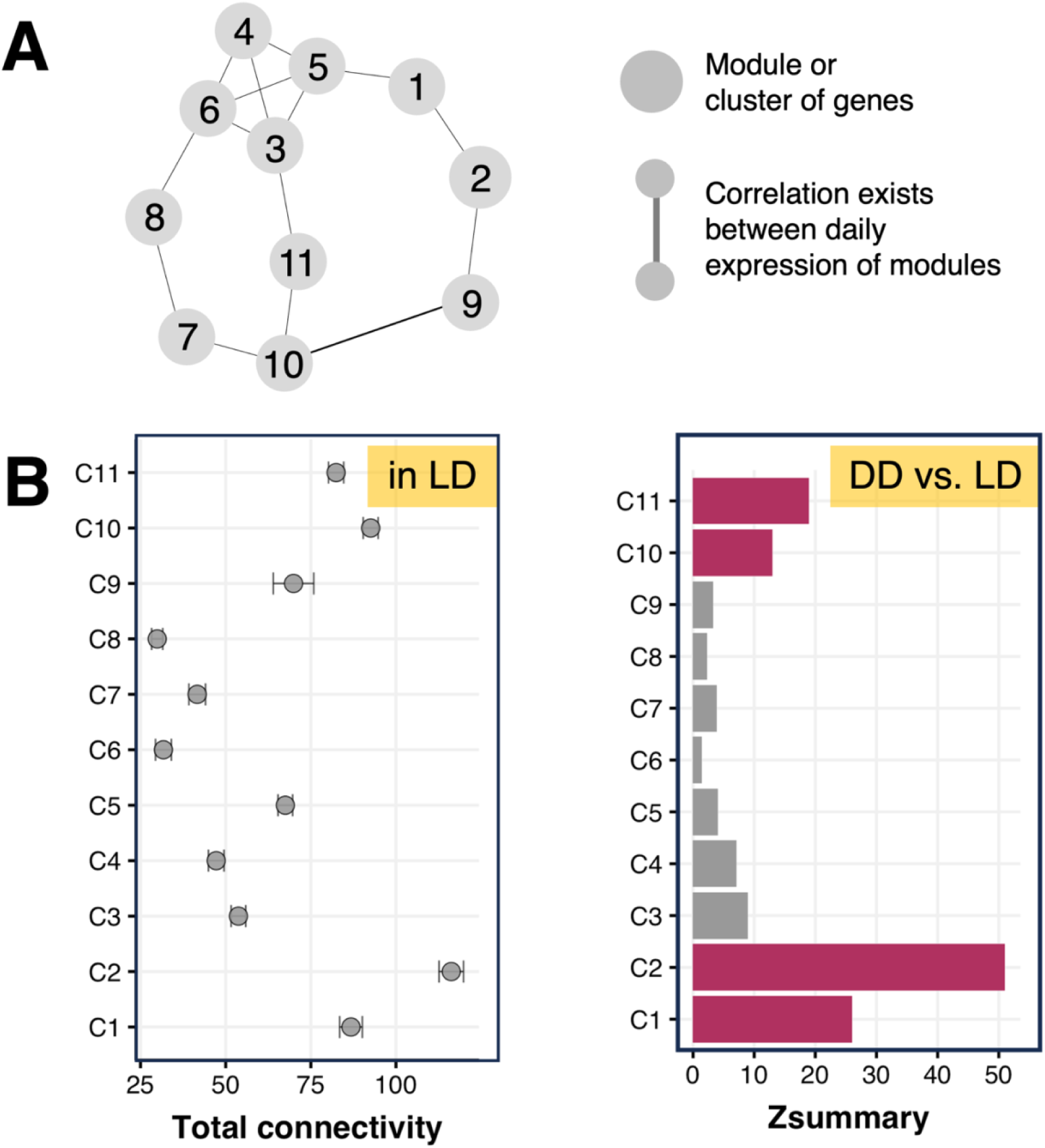
Highly connected gene modules in LD are strongly preserved in DD. (A) Model of gene co-expression network for *P. barbatus* forager brains in LD. Each node represents a module or cluster of co-expressed genes: 1 = C1, 2 = C2, and so on. Edges between two nodes represent a Kendall’s tau correlation of at least 0.6 between the two modules’ daily gene expression profiles. (B) **Left panel**: Dots represent mean total connectivity of the genes in each module; error bars represent 95% confidence intervals (see Data S3 for total connectivity of each gene). **Right panel**: The x-axis shows the Preservation Zsummary score of each module in DD vs. LD. Colored bars represent Preservation Zsummary scores above 10, corresponding to strong preservation of gene co-expression patterns.

#### Network connectivity

Four of the eleven modules showed a strong gene co-expression pattern in LD. The same four modules also showed strong evidence for preservation of their co-expression patterns in DD (Fig. 4B). A gene’s connectivity in a co-expression network measures the sum of the strengths of the edges (gene-gene correlation) connected to a gene (Langfelder & Horvath, 2008). The higher a module’s average connectivity, the more similar its patterns of expression to those of other modules. Genes in module C2 showed the highest total connectivity (116.2 ± 3.6), followed by genes in C10 (92.6± 2.2), C1 (86.8 ± 3.3), and C11 (82.4 ± 2.2) (Fig. 4B).

#### Network Preservation

The preservation (Zsummary) score for a module reflects similarity of the module’s network properties between LD and DD; a score higher than 10 indicates strong preservation in the two conditions (Langfelder *et al*., 2011). Module C2, the most highly connected module in LD, was also the most preserved one under DD vs. LD; the Zsummary preservation rankings were: C2 > C1 > C10 > C11 (Fig. 4B).

The daily expression patterns of these 4 modules of interest (C1, C2, C10, and C11) and the Pfam domains overrepresented in these modules are shown in Supplementary Fig. S6-S9. The Pfam domains enriched in each module shows the functions performed by each cluster’s translated proteins. Module C1 contains 6 overrepresented Pfam domains (Supplementary Fig. S6), including 20% of all the *P. barbatus* genes with Immunoglobulin or Immunoglobulin I-set domains that control cellular communication and immune response (Brümmendorf & Lemmon, 2001), 15% of all genes with Protein kinase domain that regulate cellular processes such as division, proliferation, apoptosis, and differentiation (Manning et al., 2002), and 18% of all genes with dual specificity protein tyrosine and serine/threonine kinases that control signal transduction, influencing almost all aspects of cellular physiology. Module C2 contains 26 overrepresented Pfam domains, including the above domains overrepresented in C1. In addition, this cluster contains more than 50% of all *P. barbatus* genes with Laminin G (cellular processes), RhoGEF (GTPase activity, signal transduction, and cytoskeletal organization), Phorbol esters/diacylglycerol binding domain or C1 domain (signal transduction), and Ligand-binding domain of nuclear hormone receptor (homeostasis, development and metabolism) domains. Module C10 is overrepresented in genes with uncharacterized protein domains and 4 known Pfam domains: THAP (DNA-binding), LSM (RNA metabolism), Helicase conserved C-terminal (RNA metabolism and DNA repair), and DEAD/DEAH box (RNA processing). Finally, module C11 is overrepresented in 8 Pfam domains: Proteasome subunit (protein quality control), WD domain, G-beta repeat (signal transduction, transcription regulation, and apoptosis), Thioredoxin (cellular redox balance), AAA+ lid (nucleotide binding and hydrolysis), RNA recognition motif (post-transcriptional gene expression), DnaJ (protein folding, assembly and translocation), Ras (cellular processes), and ATPase family (protein assembly and disassembly).

### 3. Overlap between gene co-expression networks in DD vs. LD in harvester ants

The clock-controlled genes of *P. barbatus*, ones that show 24h-rhythms in DD (*for24-Pbar-DD*), showed a significant overlap with only two of the eleven modules: C1 and C2, with stronger overlap in C2 (C2 > C1, Table 2). In comparison, *P. barbatus* genes that show 24h-rhythms in LD (*for24-Pbar-LD*) showed an overlap with four of the eleven modules; strength of overlap: C2 > C3 > C4 > C11 (Table 2). Module C2 contains most of the *P. barbatus* genes that are expressed with a 24h rhythm in both LD and DD conditions, including the gene *Per*, an essential component of the animal circadian clock.

**Table 2:** Number of overlapping genes between modules and genesets. Values indicate number of genes overlapping between a geneset of interest (columns) and each of the eleven modules of *P. barbatus* genes identified in the gene co-expression network (rows). Numbers within parenthesis indicate the odds-ratio and BH-corrected p-values (odds ratio; p-value) obtained from the Fisher’s exact test. Results from overlaps of less than 5 genes are not shown; marked as ‘–’ in the table. A p-value < 0.05 indicates significant overlap between two sets of genes, marked in bold. The number of genes in each of the eleven modules were: C1 (810), C2 (1,855), C3 (246), C4 (200), C5 (2,015), C6 (69), C7 (233), C8 (99), C9 (143), C10 (2,225), C11 (2,030).

|  | *P. barbatus*<br>genes with<br>24h rhythm<br>in LD<br>(n = 1,536) | *P. barbatus*<br>genes with<br>24h rhythm<br>in DD<br>(n = 1,547) | *P. barbatus*<br>genes positively<br>correlated<br>with reduced<br>foraging when<br>dry<br>(n = 516) | *P. barbatus*<br>genes<br>negatively<br>correlated<br>with reduced<br>foraging when<br>dry<br>(n = 516) | *C. floridanus*<br>genes with<br>24h rhythm<br>in LD<br>(n = 3,265) | *C. floridanus*<br>genes with<br>24h rhythm in<br>foragers but<br>8h rhythm in<br>nurses<br>(n = 276) |
| --- | --- | --- | --- | --- | --- | --- |
| C1 | 139<br>(1.1; 0.17) | **168**<br>**(1.5; 0)** | **134**<br>**(4.5; 0)** | — | **315**<br>**(1.3; 0)** | **59**<br>**(3.2; 0)** |
| C2 | **720**<br>**(5.7; 0)** | **510**<br>**(2.6; 0)** | **202**<br>**(3.0; 0)** | 20<br>(0.2; 1) | 633<br>(1.1; 0.4) | **101**<br>**(2.6; 0)** |
| C3 | 68<br>(2.1; 0) | 19<br>(0.5; 1) | — | 18<br>(1.5; 0.2) | 71<br>(0.8; 1) | — |
| C4 | 54<br>(2.1; 0) | 17<br>(0.5; 1) | — | 9<br>(0.9; 1) | 54<br>(0.8; 1) | — |
| C5 | 30<br>(0.1; 1) | 246<br>(0.7; 1) | 108<br>(1.0; 1) | 36<br>(0.3; 1) | 501<br>(0.6; 1) | 52<br>(0.9; 1) |
| C6 | – | – | – | – | 23<br>(1.0; 1) | – |
| C7 | 14<br>(0.3; 1) | 42<br>(1.2; 0.6) | – | 17<br>(1.5; 0.3) | 65<br>(0.8; 1) | – |
| C8 | 23<br>(1.7; 0.07) | 7<br>(0.4; 1) | – | – | 20<br>(0.5; 1) | – |
| C9 | – | 22<br>(0.98; 1) | 13<br>(1.8; 0.15) | – | 21<br>(0.4; 1) | – |
| C10 | 18<br>(0.03; 1) | 222<br>(0.5; 1) | 18<br>(0.1; 1) | **179**<br>**(1.9; 0)** | 751<br>(1.1; 0.5) | 28<br>(0.4; 1) |
| C11 | **354**<br>**(1.2; 0)** | 134<br>(0.3; 1) | 13<br>(0.1; 1) | **202**<br>**(2.7; 0)** | **765**<br>**(1.3; 0)** | 23<br>(0.4; 1) |

The clock-controlled genes in *P. barbatus* overlap with those associated in previous work with reduced foraging activity in dry conditions (Friedman *et al*., 2020). The *P. barbatus* genes that were more highly expressed in colonies that reduce foraging in dry conditions (*Pbar-rH-pos*) showed a significant overlap with two of the eleven modules, module C2 (overlap = 134 genes, odds-ratio = 4.5, p < 0.001), and module C1 (overlap = 202 genes, odds-ratio = 3, p < 0.001).

The *P. barbatus* genes that were more highly expressed in colonies that do not reduce foraging in dry conditions (*Pbar-rH-neg*) showed a significant overlap with two of the eleven modules, module C10 (overlap = 179, odds-ratio = 1.9, p < 0.001) and module C11 (overlap = 202, odds-ratio = 2.7, p < 0.001).

### 4. Comparison of rhythmically expressed genes that regulate task behavior in harvester ants and carpenter ants

There was a significant overlap between genes of *P. barbatus* and *C. floridanus* whose daily expression is significantly 24h-rhythmic under LD conditions (overlap = 567, odds-ratio = 1.5, p < 0.001). The 24h-rhythmic *C. floridanus* genes, *for24-Cflo-LD*, overlapped most strongly with *P. barbatus* modules C11 and C1. Module C11 genes show 24h-rhythms in LD but not in DD, whereas C1 genes contain the clock-controlled genes of *P. barbatus*, 24h-rhythmic in DD but not in LD (Fig. 5A; Table 2). There was no significant overlap between *for24-Cflo-LD* and module C2, which contains *P. barbatus* genes 24h-rhythmic in DD and LD (Table 2).

**Figure 5.**
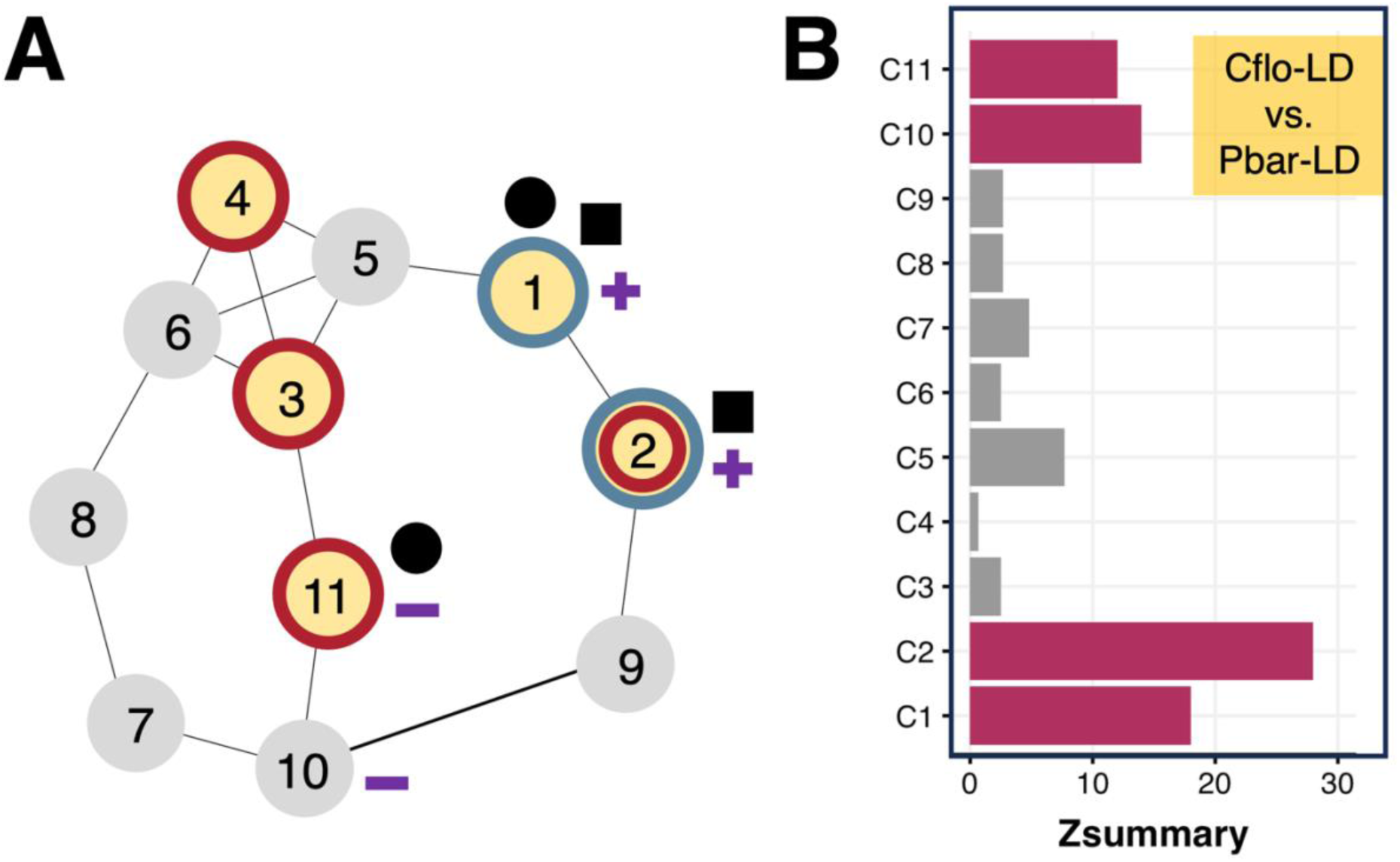
Comparison of genes associated with the circadian clock and behavioral plasticity in *P. barbatus* and *C. floridanus*. (A) Annotated GCN model in LD. Nodes and edges as in Fig. 4A. Yellow with a red circle indicates modules with 24h rhythms in LD (for24-Pbar-LD, light-driven); yellow with a blue circle indicates modules with 24h rhythms in DD (for24-Pbar-LD, clock-controlled). Symbols: **Plus**: *P. barbatus* genes whose expression is positively correlated with the extent of reduced foraging when dry (Pbar-rH-pos); **Minus**: *P. barbatus* genes whose expression is negatively correlated with the extent of reduced foraging when dry (Pbar-rH-neg); **Solid black dot**: *C. floridanus* genes expressed with 24h rhythms in LD (for24-Cflo-LD); **Solid black square**: *C. floridanus* genes with daily expression patterns linked to task (for24nur8-Cflo). (B) The x-axis shows the Preservation Zsummary score of *P. barbatus* (*Pbar-LD*) module in *C. floridanus* forager brains in LD (*Cflo-LD*). Colored bars represent Preservation Zsummary scores above 10, corresponding to strong preservation of gene co-expression patterns across the two species.

Comparing the transcriptomic network of the two species, the co-expression patterns of four of the 11 modules of *P. barbatus* genes were highly preserved in *C. floridanus*: modules C1, C2, C10, and C11 had a Z-summary score greater than 10 (Fig. 5B; preservation ranking: C2 > C1 > C10 > C11). These are the same four *P. barbatus* modules (C2, C1, C10, C11) with the highest connectivity in LD conditions, and also the most highly preserved modules in DD compared to LD (Fig. 4B).

The rhythmic genes previously linked to task in *C. floridanus* are associated with those that regulate foraging in dry conditions in *P. barbatus*. There was significant overlap between the set of 281 genes (*for24nur8-Cflo*) associated with task in *C. floridanus,* and two *P. barbatus* modules: module C1 (overlap = 59 genes, odds-ratio = 3.2, p < 0.01) and module C2 (overlap = 101 genes, odds-ratio = 2.6, p < 0.001). These two modules in *P. barbatus*, C1 and C2, contain both the clock-controlled genes of *P. barbatus* and the genes associated with the regulation of foraging in dry conditions (Fig. 5A).

## DISCUSSION

Most genes in the brains of *P. barbatus* are expressed at similar levels over a 24h day in both 12h:12h light-dark cycles (LD) and free-running, constant darkness (DD). The daily transcriptome showed that only 10 of the thousands of genes expressed in the brain show a two-fold change in average daily expression in DD compared to LD.

The number of 24h-rhythmic genes in LD (n = 1536) and DD (n = 1547) were comparable, but only 345 genes were significantly rhythmic in both LD and DD. There is similarly low overlap between genes classified as significantly 24h rhythmic in LD and DD in fruit flies (Hughes et al., 2012) and in mice (H. Li et al., 2020). In LD, daily rhythms of the transcriptome are a product of both the circadian clock and the oscillations in its environment, as in the 12:12 LD cycle. In DD, transcriptomic rhythms are mostly driven by the circadian clock. The high number of DD-only rhythmic genes indicates that these genes are under circadian control, but their rhythmic signal is weaker in LD, probably due to photic masking. The light-dark cycle, a strong zeitgeber in most species, can disrupt the rhythmic expression of these genes, either by inducing or suppressing expression regardless of the circadian clock’s phase.

The low overlap between two sets of significantly rhythmic genes has been attributed to an arbitrary choice of false discovery rate (p-value cutoffs) (Laloum & Robinson-Rechavi, 2020). To assess the quality of our daily transcriptome and rhythmicity detection algorithm, we checked five genes commonly known as housekeeping or non-cycling genes in flies and mice: *RpL32*, *RpL19*, *CLOCK*, *Gapdh*, and *EF1-α*. All five genes were not significantly 24h rhythmic in *P. barbatus* brains under LD (Supplementary Fig. S10). In DD, only *Gapdh* was classified as significantly 24h rhythmic, but the other four were not. This provides additional support that our choice of rhythmicity detection algorithm is working as expected.

To address the limitations of comparing only sets of significantly rhythmic genes, we modelled the daily transcriptome as a network, clustering genes with a similar tempo of daily expression. We found that clock-controlled genes, whose expression shows 24h rhythm in DD, form an essential node of the *P. barbatus* daily gene co-expression network (GCN). Most of the clock-controlled genes are located in two of the eleven modules of the GCN, C1 and C2.

Three separate lines of evidence indicate that module C2, the only module that shows 24h-rhythmic expression in both LD and DD, acts as an important node of the co-expression network. First, the average *P. barbatus* gene-gene connectivity in module C2 was significantly higher than all other modules in LD conditions (Fig. 4B). Second, the co-expression patterns of genes in module C2 were also most highly preserved both within *P. barbatus* (DD vs. LD in *P. barbatus*; Fig. 4B) and when compared with the distantly related species, *C. floridanus* (LD in *P. barbatus* vs. LD in *C. floridanus*; Fig. 5B). Third, module C2 contains the highly conserved clock gene *Per* and several behavior-regulating insect genes including dopamine and octopamine receptors (Fig. 3 and 5A).

In *P*. *barbatus*, *Per* shows a phase shift in free-running conditions (Fig. 3). *Per*-like phase shifts occur in other clock genes, such as the serine/threonine kinase *Nemo* identified in fruit flies (Chiu et al., 2011). There are also phase shifts, identical to those of *Per*, in neurohormone genes found in module C2 that are associated with behavior of many insect species, including *Ecdysone receptor* (*EcR*) in ants (Nemoto & Hara, 2007), *Dopamine/Ecdysteriod receptor* (*DopEcR*) in moths (Abrieux *et al*., 2014) and in ants (Das & de Bekker, 2022), and *Dopamine D2-like receptor* in flies (Lee *et al*., 2013), in ants (Friedman *et al*., 2018)); as well as neurohormone genes from C2 found in humans (*Serotonin receptor 2A* (Yeom *et al*., 2020)). More targeted methods of sequencing, such as qPCR or microarray, would be needed to determine how the tight co-expression patterns seen in Figure 3 vary among *P. barbatus* colonies, and among ant species.

Plasticity of the clock-controlled genes, in transcriptomic rhythms, seems to be associated with variation among colonies in foraging behavior (Gordon *et al*., 2023) (Figure 5). The clock-controlled genes of *P. barbatus* in modules C1 and C2 overlapped with genes previously shown to be associated with reduced foraging in dry conditions (Friedman *et al*., 2020). In that study, foragers were collected during the morning activity period, between 8 am and 12 pm, which is roughly equivalent to ZT2 (two hours after sunrise) and ZT6. Behavioral differences among colonies in the regulation of foraging (Gordon et al., 2023) may be associated with differences in the phase or amplitude of the expression of these clock-controlled genes. This variation may be important in adaptation to climate change (Gordon, 2013).

The *P. barbatus* genes that are not entirely clock-controlled, but instead require light-dark cycles for strong 24h rhythms, are also involved in the regulation of foraging in dry, hot conditions. Module C10, that shows 24h-rhythms of expression in LD but not in DD, significantly overlapped with genes associated with reduced foraging in dry conditions (Friedman *et al*., 2020) (Figure 5). Module C10, like C1 and C2, was one of the most highly connected modules of the *P. barbatus* GCN with gene-gene co-expression patterns strongly conserved in free-running DD conditions (Figure 4B). The strongly preserved co-expression patterns of C10 genes, with weakened 24h rhythms in DD, indicates that light plays a role in maintaining the 24h-rhythmic expression of this gene cluster.

Similarities in *P. barbatus* and *C. floridanus* expression patterns suggest that the links between clock-controlled genes, light-mediated rhythmic genes, and behavioral plasticity are broadly conserved across ants. The genes expressed with a daily temporal pattern under LD conditions overlapped in these two distantly related species, which differ in their chronotype (*P. barbatus*: diurnal vs. *C. floridanus*: nocturnal), diet (granivore vs. omnivore), and ecological niche (arid vs. sub-tropical). Intriguingly, the *P. barbatus* genes in modules C1 and C2, which contain most of the clock-controlled genes and the core clock gene *Per*, showed an overlap with *C. floridanus* genes whose daily rhythms are associated with task group; foragers show 24h rhythms while nurses show 8h rhythms (Das & de Bekker, 2022). Thus, both plasticity of harvester ant foragers in response to dry conditions, and of carpenter ants in task, are associated with clock-controlled genes.

A growing body of evidence links genes associated with circadian rhythms with those associated with behavioral plasticity in social insects. Plasticity in gene expression is associated with reproductive status, queen or worker, in many ant species (Morandin *et al*., 2016), as well as with task group, comparing brood care workers and foragers (Manfredini et al., 2014; Mikheyev & Linksvayer, 2015; Qiu et al., 2017). Other studies suggest an association between the daily pattern of gene expression, its phase or periodicity, and task in many ant species (Das & Gordon, 2023). Our results here, showing overlap of the clock-controlled genes previously linked to task in *C. floridanus* with those that regulate foraging activity in dry conditions in *P. barbatus,* suggest that behavioral plasticity is tied to circadian rhythms.

## ACKNOWLEDGEMENTS

We would like to thank Lauren O’Connell and her lab where some of the molecular work was done; Max Madrzyk for providing food, beverages, and company during the 24h sampling of ants; and Mila Pamplona, Mikaela Wilson, and Katie Fiocca for helpful discussions. The work was supported by grants to D.M.G. from the Templeton Foundation and the National Science Foundation (Award 1940647).

## AUTHOR CONTRIBUTIONS

- Designed and performed research: BD and DMG
- Analyzed data: BD
- Wrote the paper: BD and DMG

## STATEMENTS AND DECLARATIONS

### Ethical considerations

Not applicable

### Consent to participate

Not applicable

### Consent for publication

Not applicable

### Data availability

All cDNA sequencing data generated in this study, 24 samples of *P. barbatus* brains, will be deposited in the NCBI SRA database, and the BioProject accession number to publicly access the samples will be provided in the final manuscript. All of the code used for our analyses are publicly available on Zenodo (https://zenodo.org/doi/10.5281/zenodo.13826749). All data necessary to re-run our analyses are provided as supplementary files.

