## Supplementary Figures S1 to S10 for "Rhythmic gene expression and behavioral plasticity in harvester and carpenter ants"

1. Biology, Stanford University
2. J. Craig Venter Institute
§ corresponding author to whom proofs will be sent

### Table of Contents:

| | |
|---|---|
| Supplementary Figure **S1** | Page 2 |
| Supplementary Figure **S2** | Page 3 |
| Supplementary Figure **S3** | Page 4 |
| Supplementary Figure **S4** | Page 5 |
| Supplementary Figure **S5** | Page 6 |
| Supplementary Figure **S6 – S9** | Page 7 - 10 |
| Supplementary Figure **S10** | Page 11 |

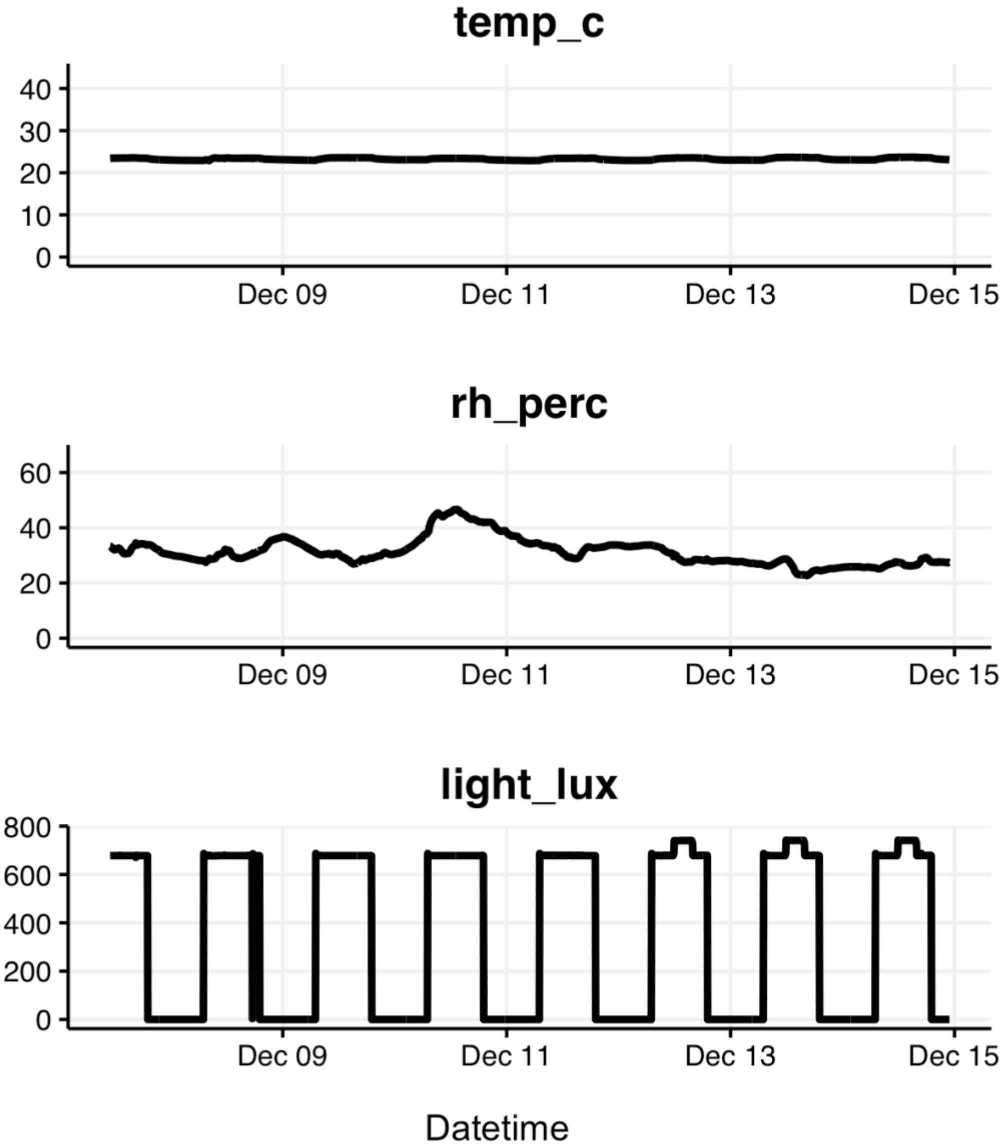


**Supplementary Figure S1:** Environmental conditions during the seven-day entrainment period under 12h:12h LD conditions. The y-axis shows (**top**) temperature in degree Celsius, (**middle**) relative humidity in percentage relative humidity, and (**bottom**) light levels in lux.

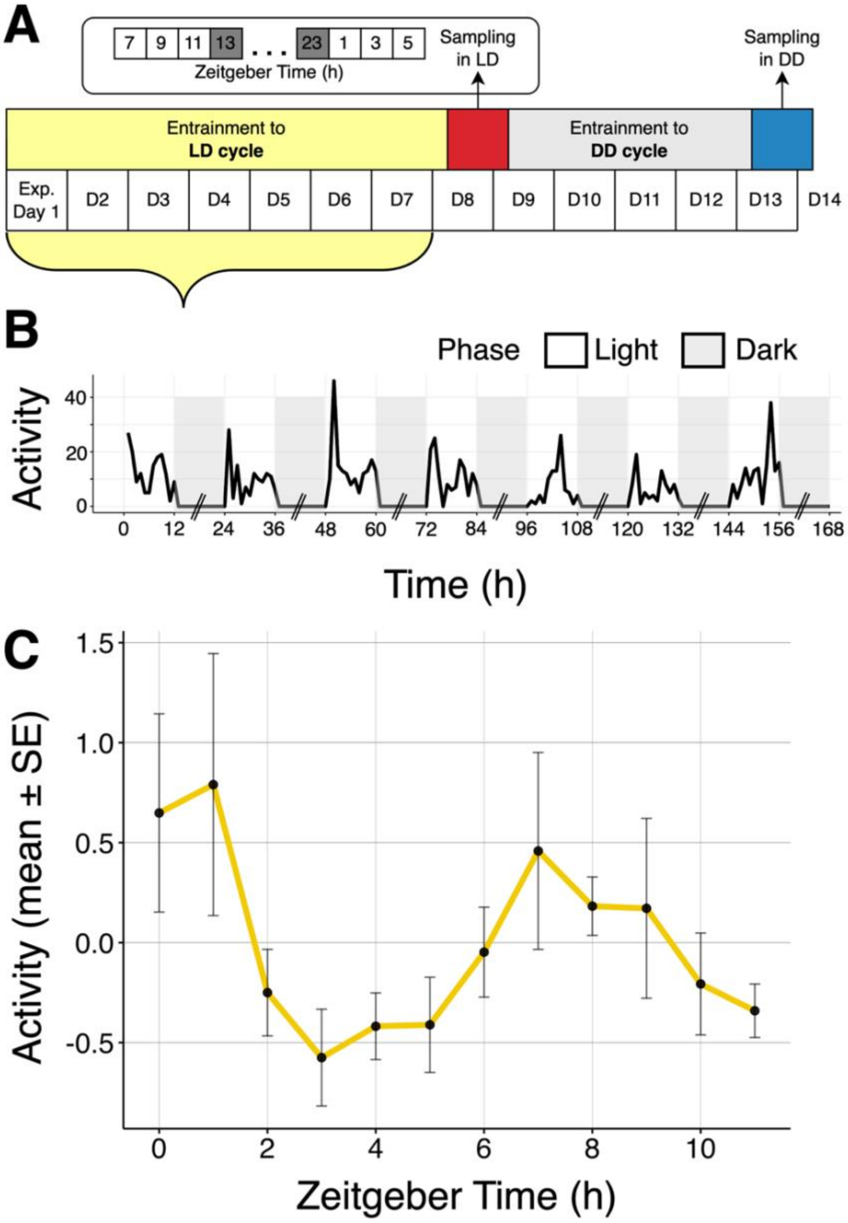


**Supplementary Figure S2**: (A) Experimental design; LD = 12:12h light-dark cycles, DD = constant darkness. (B) The y-axis shows total activity, the sum of number of ants in the foraging arena in 6 counts made over 3 min, during the 7 days of entrainment to LD cycle. The x-axis shows time since the start of the experiment. The light phases began at 0h, 24h, 48h, and so on; dark phases began at 12h, 36h, 60h, and so on. Values during the dark phase are shown as zero when activity was not measured. (C) Average daytime fluctuations in total activity. The y-axis shows mean (± SE) activity in the foraging arena during the daytime over 7 days of entrainment under 12:12 LD cycle. Zeitgeber time = time since the lights were on. Assessment of rhythmic day-time activity was performed using Wavelet decomposition methods, see "Monitoring colony activity rhythm" in Materials and Methods.

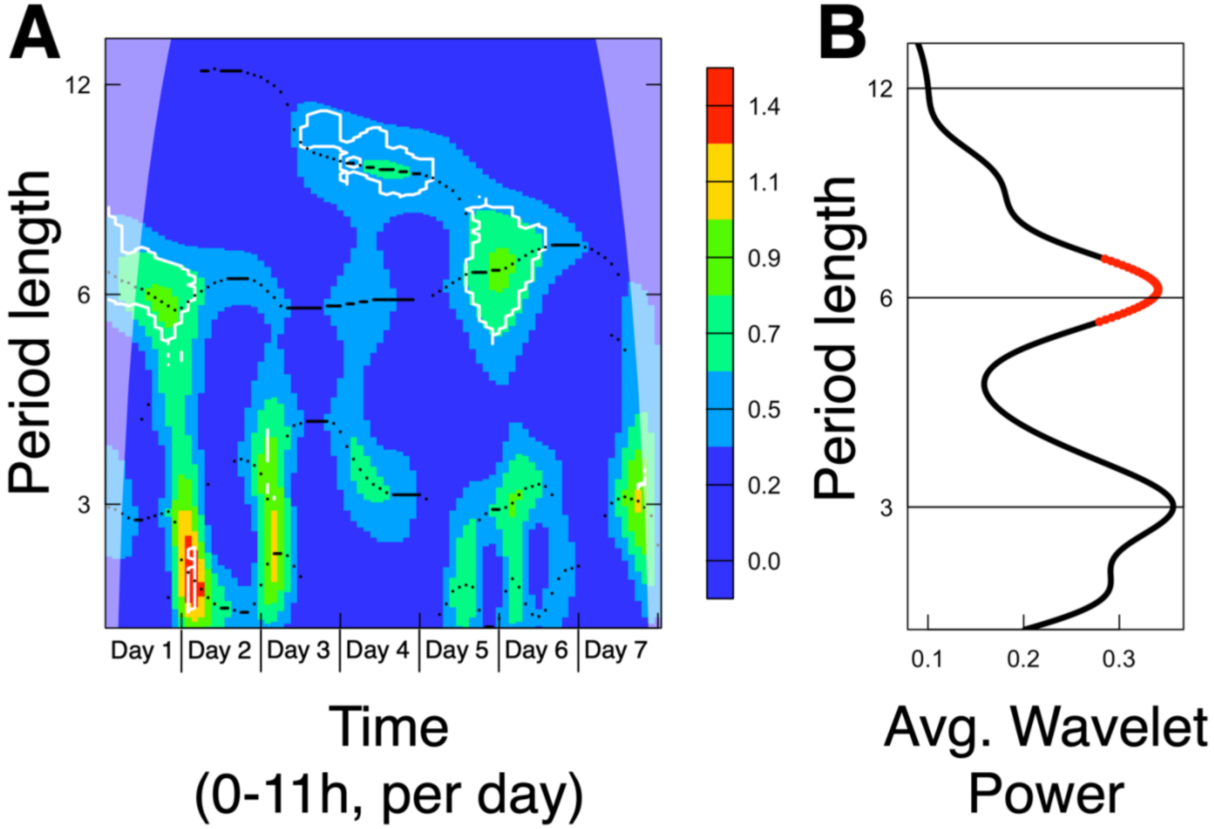


**Supplementary Figure S3:** (A) Wavelet power spectrum for total activity during the light phase over 7 consecutive days. The x-axis shows concatenated time (in hours) across the seven days of LD entrainment, during the light phase only (0-11h in the graph = 0-11h on day one of the experiment, 11-23h in the graph = 0-11h on day two, and so on). The y-axis shows the Fourier periods. Colors indicate power, how much a given periodicity contributes to the fluctuations in observed timeseries. (B) Average power for the wavelet power spectra shown in A. The y-axis is the same as in (A), and the x-axis shows the average power calculated from the wavelet power spectra in (A). Red dots highlight the periodicities that have a significant average wavelet power ($p < 0.05$).

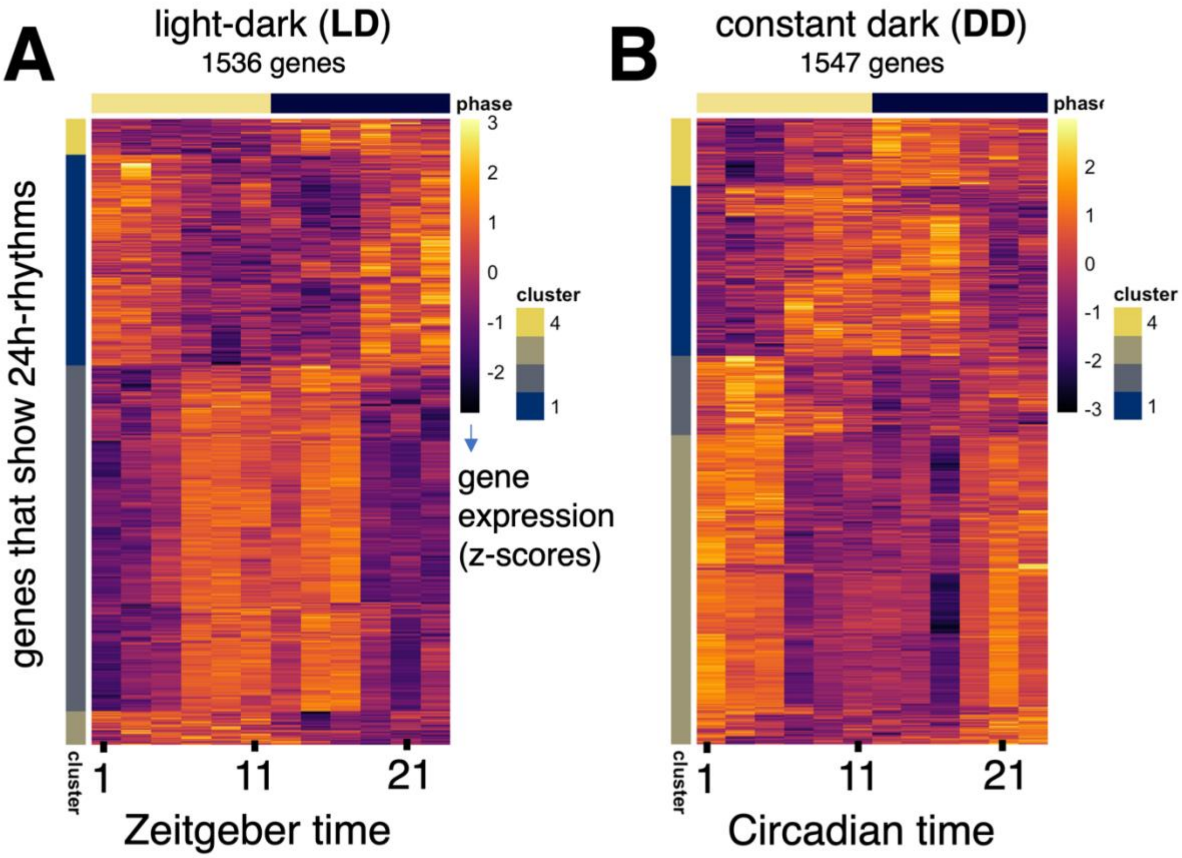


**Supplementary Figure S4:** Heatmap of daily expression (z-score) patterns of (**A**) the 1536 genes expressed with a 24h-rhythm in *P. barbatus* brains in LD and (**B**) the 1547 genes expressed with a 24h-rhythm in *P. barbatus* brains in DD conditions. Each row represents a single gene, and each column represents the time of day at each 2 hr interval at which a sample was collected, shown in chronological order from left to right. Colors indicate normalized gene expression values (z-scores): negative numbers (dark colors) indicate lower gene expression relative to their daily average, and positive numbers (lighter colors) indicate higher gene expression.

| Sample | Qubit BR-RNA $C_i$ (ng/uL) | vol_RNA actually used (µL) | qty_RNA actually used (ng) | NEB Index primer used | Library conc. Qubit HS (ng/uL) | Vol_library left after Qubit & Tapestation | Libraries prepped on | fragment size of dsDNA (bp) |
|---|---|---|---|---|---|---|---|---|
| LD-07 | 73.2 | 6.16 | 450.91 | S1-01 | 2.66 | 18 | 17-Jan-23 | 270 |
| LD-09 | 69.6 | 6.48 | 451.01 | S1-07 | 1.96 | 18 | 20-Jan-23 | 282 |
| LD-11 | 71 | 6.34 | 450.14 | S1-02 | 1.33 | 18 | 17-Jan-23 | 272 |
| LD-13 | 56.6 | 7.96 | 450.54 | S1-08 | 6.36 | 18 | 20-Jan-23 | 310 |
| LD-15 | 56.8 | 7.94 | 450.99 | S1-03 | 0.24 | 18 | 17-Jan-23 | 269 |
| LD-17 | 49.6 | 9.08 | 450.37 | S1-09 | 7.84 | 18 | 20-Jan-23 | 303 |
| LD-19 | 60.2 | 7.48 | 450.30 | S1-04 | 4.94 | 18 | 17-Jan-23 | 303 |
| LD-21 | 54.6 | 8.26 | 451.00 | S1-10 | 17 | 18 | 20-Jan-23 | 303 |
| LD-23 | 63.6 | 7.08 | 450.29 | S1-05 | 0.18 | 18 | 17-Jan-23 | 300 |
| LD-01 | 68.8 | 6.56 | 451.33 | S1-11 | 11.5 | 18 | 20-Jan-23 | 302 |
| LD-03 | 67.4 | 6.68 | 450.23 | S1-06 | 1.64 | 18 | 17-Jan-23 | 318 |
| LD-05 | 67.4 | 6.68 | 450.23 | S1-12 | 10.3 | 18 | 20-Jan-23 | 298 |
| DD-07 | 58.8 | 7.66 | 450.41 | S2-13 | 2.02 | 18 | 24-Jan-23 | 278 |
| DD-09 | 60 | 7.5 | 450.00 | S2-20 | 11.2 | 18 | 26-Jan-23 | 312 |
| DD-11 | 59.6 | 7.56 | 450.58 | S2-14 | 7.72 | 18 | 24-Jan-23 | 322 |
| DD-13 | 63.6 | 7.08 | 450.29 | S2-21 | 16.1 | 18 | 26-Jan-23 | 307 |
| DD-15 | 75.6 | 5.96 | 450.58 | S2-15 | 9.58 | 18 | 24-Jan-23 | 325 |
| DD-17 | 65.2 | 6.92 | 451.18 | S2-22 | 25 | 18 | 26-Jan-23 | 326 |
| DD-19 | 69.8 | 6.46 | 450.91 | S2-16 | 13.4 | 18 | 24-Jan-23 | 317 |
| DD-21 | 67.8 | 6.64 | 450.19 | S2-23 | 16.9 | 18 | 26-Jan-23 | 307 |
| DD-23 | 64.6 | 6.98 | 450.91 | S2-18 | 17.2 | 18 | 24-Jan-23 | 294 |
| DD-01 | 65 | 6.94 | 451.10 | S2-25 | 18 | 18 | 26-Jan-23 | 317 |
| DD-03 | 72.8 | 6.2 | 451.36 | S2-19 | 7.96 | 18 | 24-Jan-23 | 323 |
| DD-05 | 59.2 | 7.62 | 451.10 | S2-27 | 20 | 18 | 26-Jan-23 | 307 |

**Supplementary Figure S5:** Sample quality of RNA extraction and DNA library preps. The table columns have the following meanings. **Sample**: light-dark cycle and timepoint of collection, **Qubit BR-RNA**: concentration of mRNA extracted from three ant brains pooled together, **vol_RNA**: volume of mRNA collected, **qty_RNA**: starting amount of mRNA (in ng) used for cDNA library preps, **NEB index**: ID of the unique primer used for sequencing the cDNA, **Library conc.**: concentration of the cDNA fragments created from the starting mRNA, **Vol_library**: volume of cDNA libraries available for sequencing, **Libraries prepped on**: date on which cDNA libraries were generated, **Fragment size**: average fragment size (in bp) of the resulting cDNA libraries. Qubit plots available upon request.

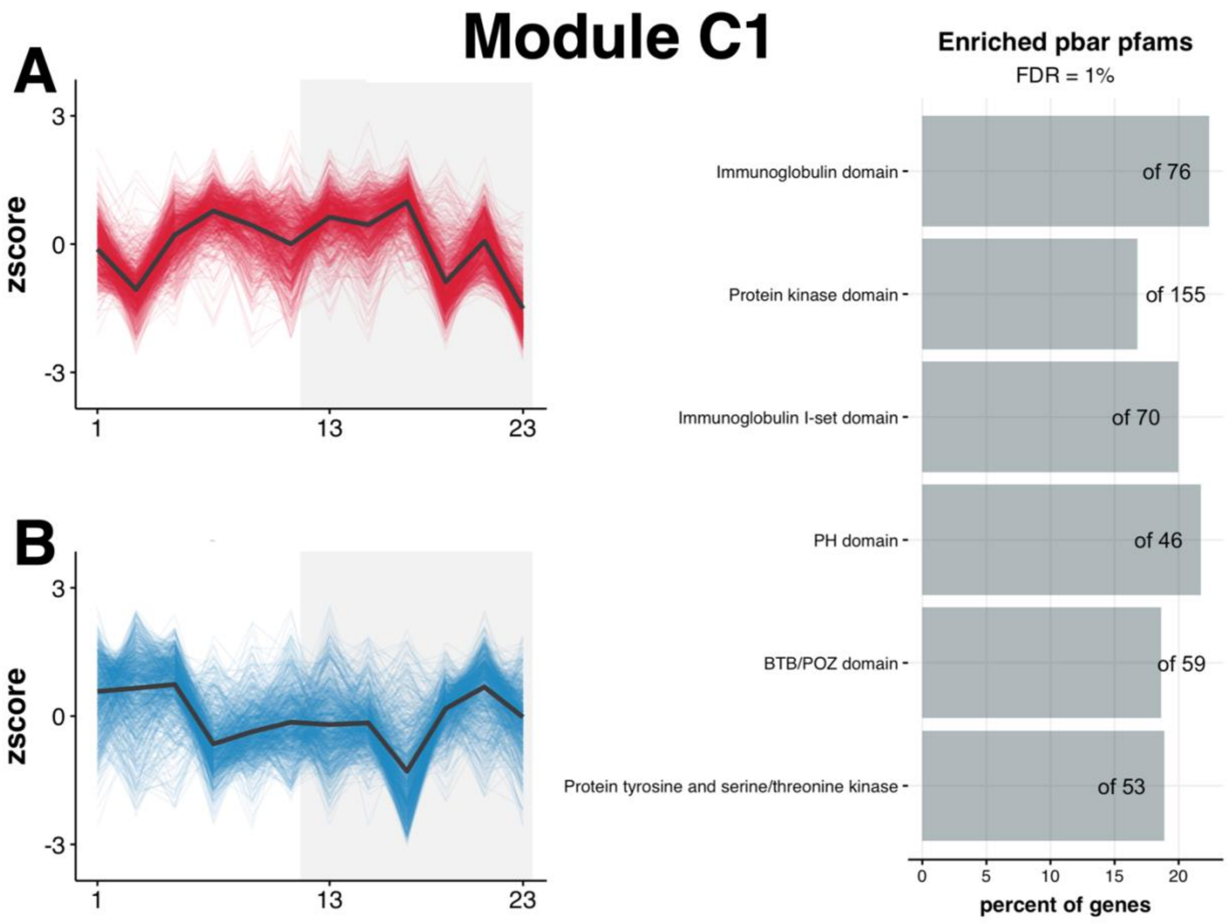


**Supplementary Figure S6:** Daily tempo of gene expression in module C1 and the Pfam domains overrepresented in these genes (n = 810). The figure on the left shows the daily expression of all genes in module C1 in (A) LD conditions and (B) DD conditions. The y-axis shows amount of gene expression (z-scored FPKM values), and the x-axis shows time in hours (ZT in LD and CT in DD). The shaded portion indicates nighttime in LD (ZT12 to ZT24) and subjective nighttime in DD (CT12 to CT24). The figure on the right shows the overrepresented Pfam domains identified using a hypergeometric test using a false discovery rate of 1%, including only Pfam terms that are present in at least 10 genes in *P. barbatus*. The numbers displayed on top of the bars show the number of genes that have the Pfam term in the entire *P. barbatus* genome. The x-axis shows the percent of all genes annotated with the Pfam domain that are present in the module.

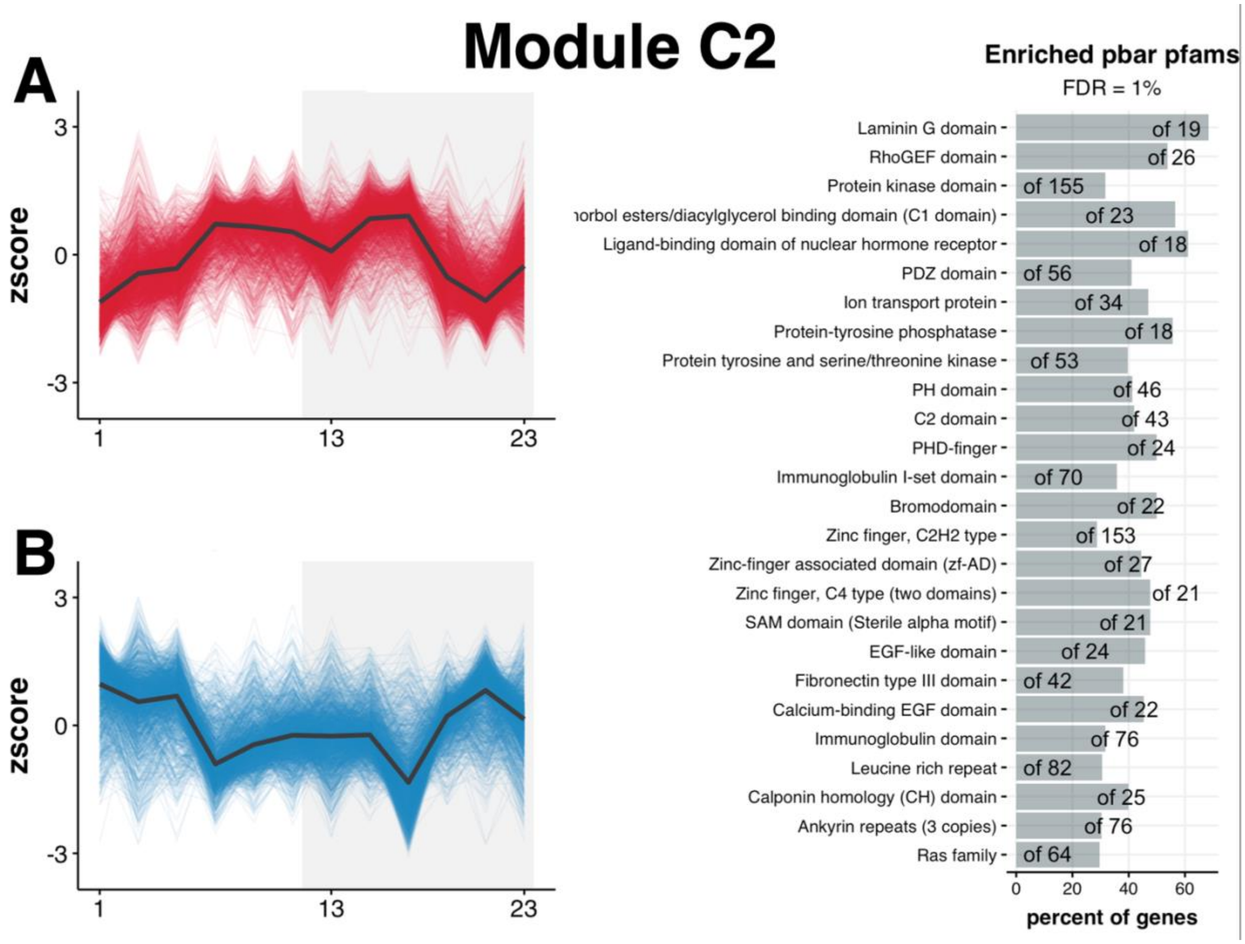


**Supplementary Figure S7:** Daily tempo of gene expression in module C2 and the Pfam domains overrepresented in these genes (n = 1855). The figure on the left shows the daily expression of all genes in module C1 in (A) LD conditions and (B) DD conditions. The y-axis shows amount of gene expression (z-scored FPKM values), and the x-axis shows time in hours (ZT in LD and CT in DD). The shaded portion indicates nighttime in LD (ZT12 to ZT24) and subjective nighttime in DD (CT12 to CT24). The figure on the right shows the overrepresented Pfam domains identified using a hypergeometric test using a false discovery rate of 1% and only including Pfam terms that are present in at least 10 genes in *P. barbatus*. The numbers displayed on top of the bars show the number of genes that have the Pfam term in the entire *P. barbatus* genome. The x-axis shows the percent of all genes annotated with the Pfam domain that are present in the module.

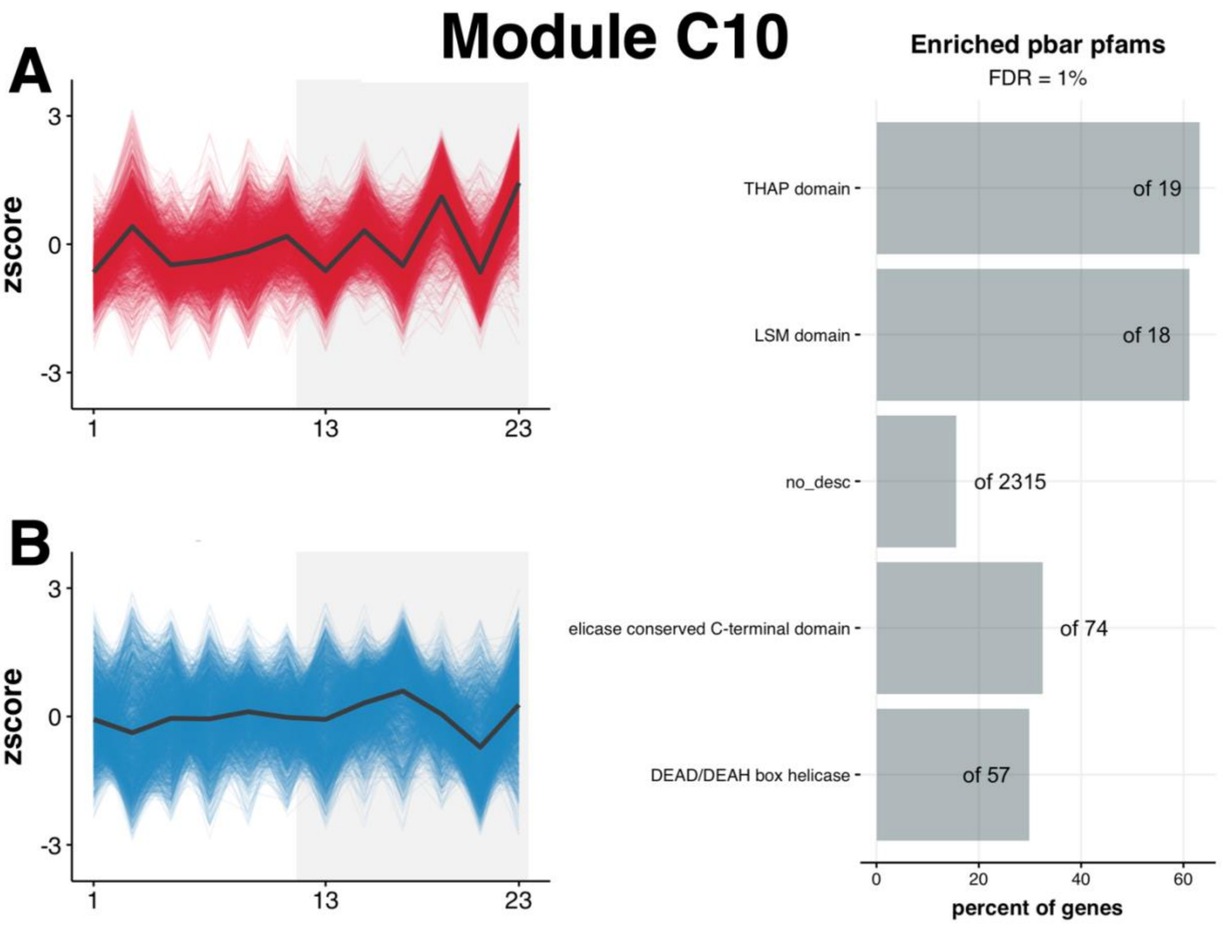


**Supplementary Figure S8:** Daily tempo of gene expression in module C10 and the Pfam domains overrepresented in these genes (n = 2225). The figure on the left shows the daily expression of all genes in module C1 in (A) LD conditions and (B) DD conditions. The y-axis shows amount of gene expression (z-scored FPKM values), and the x-axis shows time in hours (ZT in LD and CT in DD). The shaded portion indicates nighttime in LD (ZT12 to ZT24) and subjective nighttime in DD (CT12 to CT24). The figure on the right shows the overrepresented Pfam domains identified using a hypergeometric test using a false discovery rate of 1% and only including Pfam terms that are present in at least 10 genes in *P. barbatus*. The numbers displayed on top of the bars show the number of genes that have the Pfam term in the entire *P. barbatus* genome. The x-axis shows the percent of all genes annotated with the Pfam domain that are present in the module.

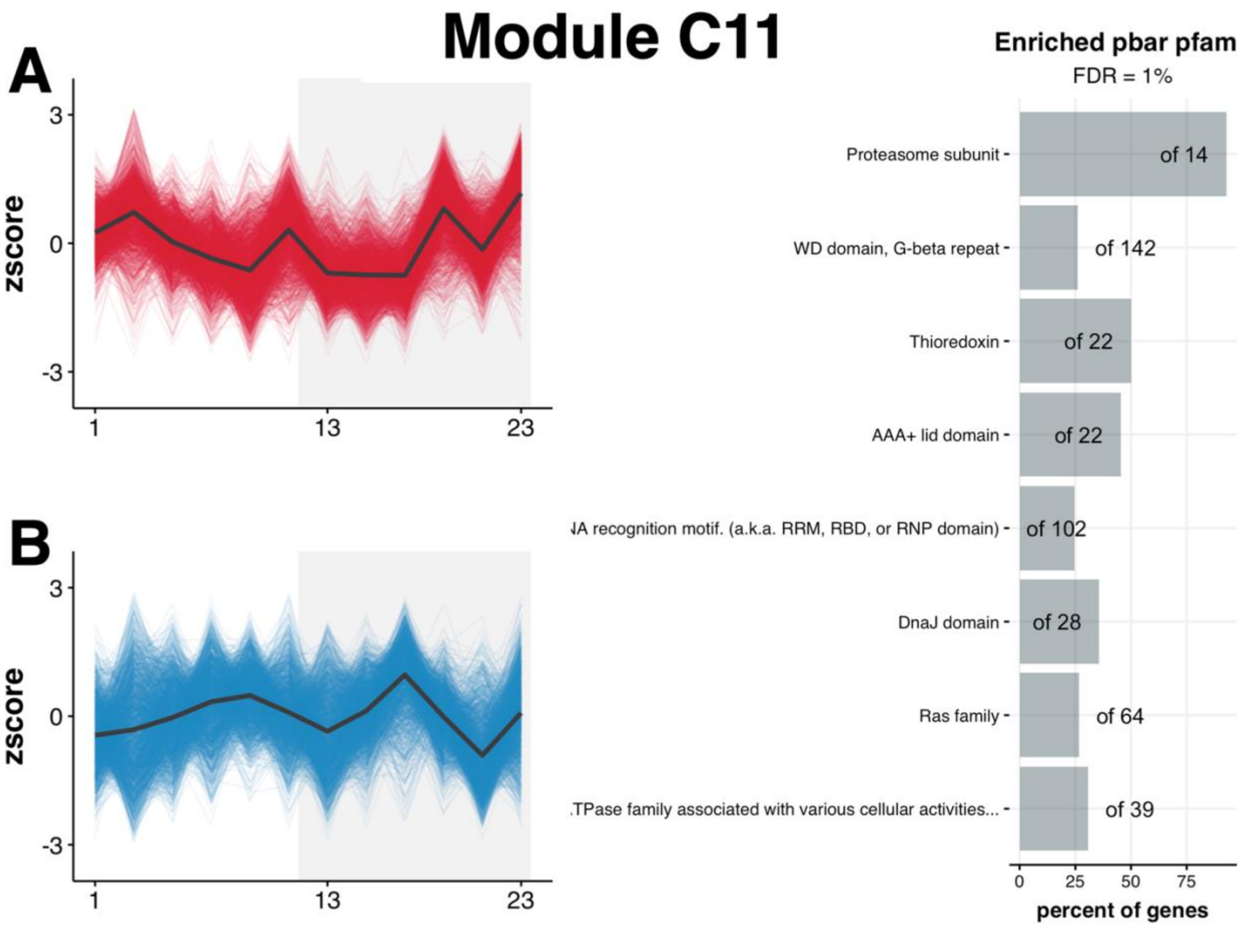


**Supplementary Figure S9:** Daily tempo of gene expression in module C11 and the Pfam domains overrepresented in these genes (n = 2030). The figure on the left shows the daily expression of all genes in module C1 in (A) LD conditions and (B) DD conditions. The y-axis shows amount of gene expression (z-scored FPKM values), and the x-axis shows time in hours (ZT in LD and CT in DD). The shaded portion indicates nighttime in LD (ZT12 to ZT24) and subjective nighttime in DD (CT12 to CT24). The figure on the right shows the overrepresented Pfam domains identified using a hypergeometric test using a false discovery rate of 1% and only including Pfam terms that are present in at least 10 genes in *P. barbatus*. The numbers displayed on top of the bars show the number of genes that have the Pfam term in the entire *P. barbatus* genome. The x-axis shows the percent of all genes annotated with the Pfam domain that are present in the module.

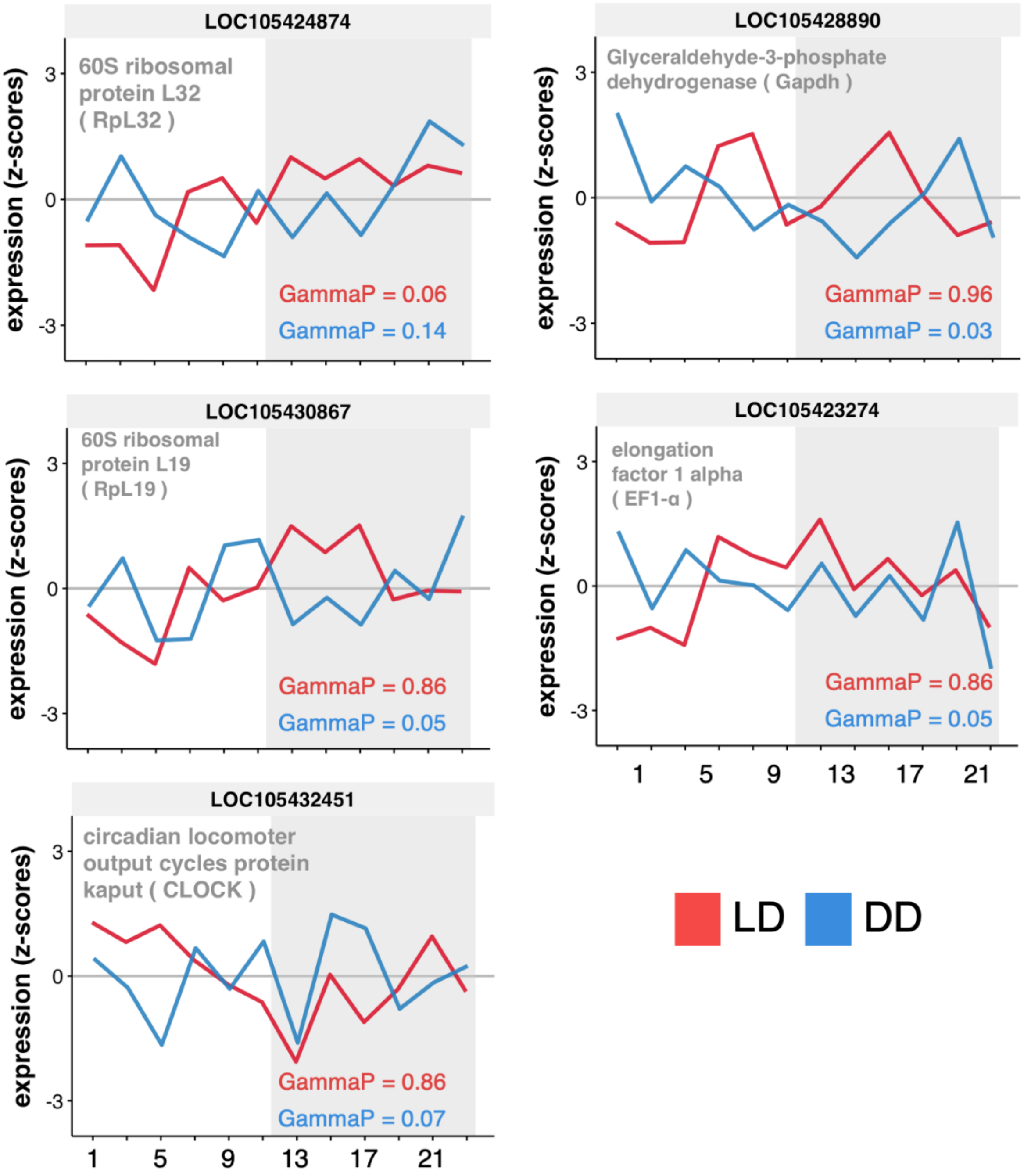


**Supplementary Figure S10:** Daily tempo of gene expression in *P. barbatus* brains for five commonly known housekeeping or non-cycling genes in LD (red) and DD (blue) conditions. For each of the genes' daily tempo, in LD or DD, the GammaP values obtained from running eJTK cycle to test for 24h rhythms are shown. Significant 24h rhythm is inferred when GammaP is less than 0.05.
